# Unmasking Early Hepatorenal Fibrosis in Polycystic Kidney Disease Using Noninvasive Precision Molecular MRI of Collagen

**DOI:** 10.64898/2026.07.19.739445

**Authors:** Oluwabukola S. Bamishaye, Francis Akinlotan, Phillip H. Chumley, Jingjuan Qiao, Xiu Chen, Farzaneh Dorabadizare, Roye Ronald, Annye Bennett, Mandy J. Croyle, Khan Hekmatyar, Alton B. Farris, Anna G. Sorace, Harrison Kim, Jian-Xiong Wang, Hans E. Grossniklaus, Bradley K. Yoder, Michal Mrug, Jenny J. Yang

## Abstract

Fibrosis begins early in autosomal dominant polycystic kidney disease (ADPKD) and drives its progression. Collagen deposition has been observed to increase around cysts in human ADPKD kidneys and livers. Currently, no clinical methods are available to detect early fibrosis before irreversible structural and functional decline. Here, we report fibrosis imaging using Gd-hProCA32.Collagen, a collagen-targeting protein MRI contrast agent for precision molecular MRI (pMRI). Gd-hProCA32.Collagen binds collagen type I with micromolar affinity, exhibiting higher relaxivity than clinical agents at 1.4 T, blood clearance within 72 hours, and no evident acute toxicity. To assess the ability of Gd-hProCA32.Collagen to detect ADPKD-related fibrosis, renal collagen deposition was increased by 11 weeks of age in *Pkhd1*^PCK/PCK^ (PCK) rats. At 26 weeks, Gd-hProCA32.Collagen, given at a 10-fold lower dose than gadobutrol, produced sustained kidney and liver enhancement through 48 hours, whereas enhancement washed out in wild-type organs and with a non-targeted control protein. Gd-hProCA32.Collagen also increased the number of detected renal cysts per animal by 25% compared with 3% for gadobutrol (*P* < 0.0001), mainly small and medium cysts outlined by pericystic collagen, and revealed striation-like enhancement at microcyst clusters. Regional enhancement correlated with histological collagen (*r* = 0.72), whereas gadobutrol enhancement did not (*r* = 0.21). Kidney and liver enhancement was also observed in a *Pkd2* mutant mouse model. These findings reveal a previously inaccessible subclinical fibrotic phase of polycystic kidney and liver disease and support collagen-targeted pMRI for noninvasive detection and mapping of fibrosis when conventional biomarkers remain uninformative.

**One-sentence summary:** We report a first-in-class collagen-targeted MRI contrast agent with Precision molecular MRI that noninvasively reveals a previously inaccessible subclinical fibrotic phase of polycystic kidney disease across kidney and liver, overcoming limitations of current diagnostic methodology.

## Introduction

Chronic kidney disease (CKD) encompasses a spectrum of progressive disorders that impair renal structure and function (*1*). CKD affects over 35 million Americans (∼14% of adults), and most cases remain undetected until advanced stages, underscoring a critical gap in early diagnosis, risk assessment, and intervention. The landmark 2002 classification guidelines marked a pivotal shift in recognizing CKD as a global public health concern and advocated early-stage management by general practitioners and internists (*1*). Early detection is vital since timely intervention can slow progression, reduce complication risk, and improve long-term outcomes (*2*).

The leading inherited cause of end-stage kidney disease (ESKD), and the fourth leading ESKD cause overall, is Autosomal Dominant Polycystic Kidney Disease (ADPKD). ADPKD is characterized by progressive renal and liver cyst expansion, interstitial fibrosis, and eventual loss of renal function (*3, 4*) (Fig. 1). Slowing cyst growth and fibrosis thus forms the critical foundation for future ADPKD care (*5–8*). Comprehensive risk stratification with longitudinal follow-up is essential for identifying patients at risk, selecting appropriate management strategies, and monitoring therapeutic responses (*9, 10*).

**Fig. 1.**
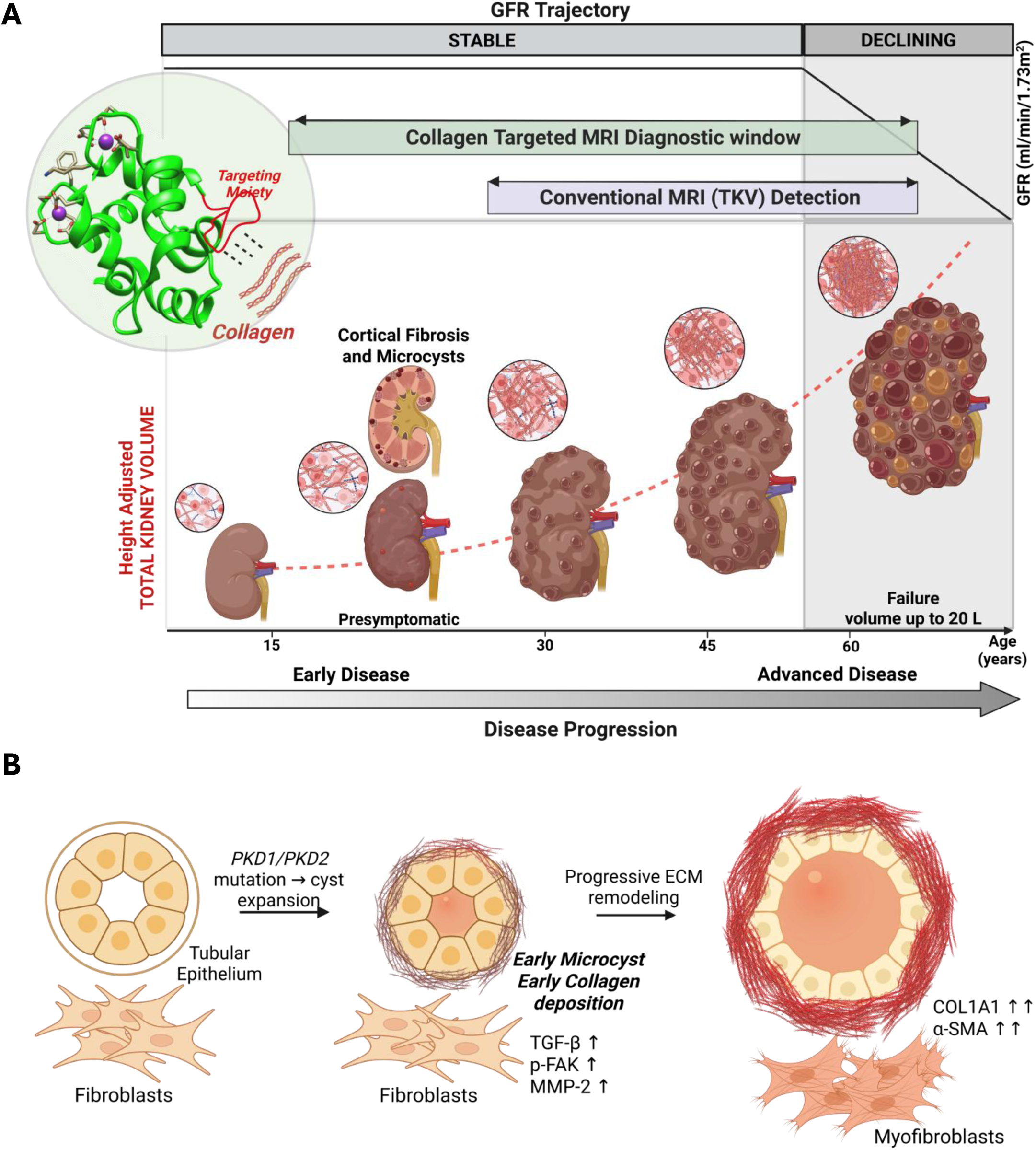
The role of collagen-targeted precision MRI in early detection of renal fibrosis during the pre-symptomatic window of ADPKD progression. **(A)** Schematic depicting the relationship between glomerular filtration rate (GFR) trajectory and height-adjusted total kidney volume (htTKV) in ADPKD progression. Kidney cross-sections illustrate progressive cystic and fibrotic burden from early pre-symptomatic disease (near normal volume in early age) through advancing disease and ensuing function impairment (increased kidney volume by middle age), and ESKD (large kidney volume in more advanced ages). GFR remains stable for decades despite ongoing structural injury, declining only in advanced disease, while htTKV (red dashed line) progressively grows beyond the functionally silent phase. Schematic insets illustrate progressive cortical fibrosis and microcyst formation culminating in near-total replacement of functional nephrons by collagen-rich cysts. The diagnostic window for collagen-targeted MRI may substantially precede the detection threshold of conventional MRI. Gd-hProCA32.Collagen comprises a protein scaffold (green ribbon helix) combined with a collagen-targeting moiety (red ribbon loop) conjugated to gadolinium ions (purple spheres) that selectively binds extracellular collagen type I (red structure), the principal matrix component of renal fibrosis in ADPKD. **(B)** Schematic of pericystic fibrosis progression at the cellular level. *PKD1* or *PKD2* mutation drives tubular epithelial cyst expansion with early pericystic collagen deposition and activation of pro-fibrotic signaling, fibroblast-to-myofibroblast differentiation, and progressive extracellular matrix remodeling characterized by upregulation of collagen type I (COL1A1) expression. *(A) Created in BioRender. Yang lab, J.* (*2026*) https://BioRender.com/zmrnlkw *(B) Created in BioRender. Yang lab, J.* (*2026*) https://BioRender.com/snqle4e

Renal fibrosis is among the central drivers of irreparable renal function loss common to most forms of CKDs, including ADPKD (*11*). Fibrosis is defined by remodeling of the extracellular matrix (ECM) that involves excessive type I collagen deposition. This leads to progressive disruption of renal architecture well before measurable functional decline (*12–16*). The consequences include irreversible loss of renal parenchyma, functional nephron depletion, and ultimately ESKD requiring dialysis or transplantation (*17, 18*). The pathogenesis of renal fibrosis is complex, attributed in part to persistent activation of myofibroblasts and ensuing abnormal ECM remodeling, resulting in progressive architectural distortion and irreversible loss of functional renal parenchyma (*19–25*). Notably, interstitial fibrosis is now recognized as an early and active disease driver rather than a late consequence of cyst expansion, preceding, accompanying, and potentially accelerating cystogenesis (*26, 27*) (Fig. 1). In addition to the renal involvement, hepatic manifestations affect more than 90% of ADPKD patients older than 35 years (*28*). The hepatic manifestations may involve cholangiocyte proliferation, ductal plate malfunction, and progressive fibrosis, all of which may promote the formation of a collagen-rich microenvironment analogous to that observed in ADPKD kidneys (*29–31*).

Despite its central role in ADPKD progression and prediction of renal outcomes (*14–16, 32*), renal fibrosis cannot be directly assessed because a kidney biopsy is not practical for routine longitudinal assessment, and is even contraindicated in ADPKD ^32^. Therefore, the burden of renal fibrosis remains virtually unknown from early ADPKD until conventional renal function markers deteriorate from normal ranges due to substantial structural damage and functional tissue loss (*3, 33, 34*). In very early or mild disease, even the cyst volume-driven total kidney volume (TKV; often derived from T2-weighted MRI) may remain indiscriminatory, creating a diagnostic and prognostic blind spot. Intricacies of fibrosis assessment in cystic livers resemble those of cystic kidneys.

MRI has emerged as a promising noninvasive diagnostic and prognostic modality for ADPKD due to its favorable safety profile (e.g., lack of ionizing radiation), high soft-tissue contrast, and capability for multiparametric assessment (*35–37*). In particular, it has become the preferred method for assessment of TKV, a foundation of broadly used imaging classifications for risk prediction assessment (*3*) (*38–40*). However, with respect to fibrosis detection, both routine and advanced MRI techniques (e.g., diffusion MRI, magnetization transfer, arterial spin labeling, BOLD MRI, and MR elastography) have insufficient molecular specificity for fibrosis and suffer from technical variability and lack of quantitative biomarker sensitivity (*38, 40–45*). Even these techniques cannot accurately detect microcysts (cysts <2–7 mm, MRI resolution limit) that likely contribute disproportionately to the ADPKD progression conductive microenvironment, given their overwhelming numerical predominance. Notably, smaller cysts have a disproportionately larger epithelial surface area relative to volume, conferring greater pro-fibrotic potential to promote abnormal ECM remodeling and function loss (*46*). Even with current MRI methods, total cyst number and cyst parenchyma surface area offer superior prediction of the slope of kidney function decline and progression to kidney failure compared to TKV (*47*). In addition, the dynamics of cyst volume changes over time are heterogeneous, with some cysts expanding rapidly, others remaining stable, and some regressing, likely leaving fibrotic remnants (*48, 49*). With current MRI techniques, detecting these processes remains technically challenging, clinically impractical, or both. Imaging approaches capable of directly detecting and quantifying microcysts and early-stage multi-organ fibrosis thus remain a major unmet need.

Current gadolinium-based contrast agents (GBCAs) have low relaxivity (r₁ and r₂ < 10 mM⁻¹s⁻¹), requiring the use of high doses for modest signal enhancement, raising potential safety concerns. Although newer macrocyclic GBCAs exhibit improved stability, current FDA, ACR, and NKF guidelines still recommend caution in patients with CKD, including ADPKD (*50–54*). Emerging evidence further suggests that repeated renal injury episodes may accelerate fibrosis progression and cyst-associated remodeling in PKD kidneys (*53, 55–57*). Because ADPKD kidneys exhibit altered tubule architecture, impaired flow dynamics, regional hypoxia, mitochondrial dysfunction, and increased susceptibility to injury (*58–62*), even transient contrast-mediated tubular stress may contribute to disease progression. Critically, these agents lack molecular specificity and cannot target collagen, precluding quantitative assessment of fibrosis (*63–68*). Efforts to develop targeted contrast agents by modifying small-molecule chelates have been limited by trade-offs between metal stability, relaxivity, and target binding, as well as limited expression of molecular biomarkers such as receptors and collagen (in the sub-nM to µM range) (*37, 69–77*).

To overcome these limitations, we developed Gd-hProCA32.Collagen, a type I collagen-targeted, protein-based MRI contrast agent that enables direct molecular imaging of extracellular matrix remodeling. Building on our prior work in engineering high-relaxivity protein contrast agents (*78–80*), we hypothesized that collagen-targeted precision molecular MRI (pMRI) would enable sensitive detection of early fibrosis at substantially lower contrast-agent doses than conventional MRI. We tested whether this approach could reveal subclinical renal fibrosis in experimental polycystic kidney disease, whether MRI enhancement correlated quantitatively with histological fibrosis, and whether it could spatially resolve early cortical fibrotic remodeling when total kidney volume (TKV) and standard clinical biomarkers remained unchanged. We further evaluated its ability to detect and characterize hepatic fibrosis noninvasively, extending the approach to renal and hepatic assessment within a single imaging framework. These studies establish the preclinical basis for evaluating collagen-targeted pMRI as a tool for early disease assessment and longitudinal monitoring, with future studies needed to determine its utility for risk stratification, outcome prediction, and assessment of therapeutic response in ADPKD.

## Results

### Fibrillar collagen deposition is increased in human ADPKD kidney and polycystic liver tissue

To validate the clinical relevance of collagen as an imaging target, we quantified collagen proportionate area (CPA) in Picrosirius red–stained sections of human ADPKD kidneys, PLD livers, and normal controls. Collagen deposition in ADPKD kidneys was concentrated in pericystic and interstitial regions, and CPA was about threefold higher than in normal kidneys (34.0 ± 4.9% versus 9.9 ± 1.1%; *n* = 5 per group; *P* < 0.0001) (Fig. 2A). The pronounced enrichment of collagen, especially in areas surrounding cysts, indicated active fibrotic remodeling associated with disease progression. Parallel analysis of PLD livers revealed similarly elevated collagen deposition, with intense Sirius Red staining observed in the pericystic regions compared to normal liver tissue, with intense pericystic staining and higher CPA than normal liver (21.5 ± 4.8% versus 7.6 ± 1.1%; *n* = 5; *P* < 0.001) (Fig. 2B). These findings confirmed that fibrosis and its enrichment in pericystic regions are a shared pathological feature across renal and hepatic ADPKD manifestations.

**Fig. 2.**
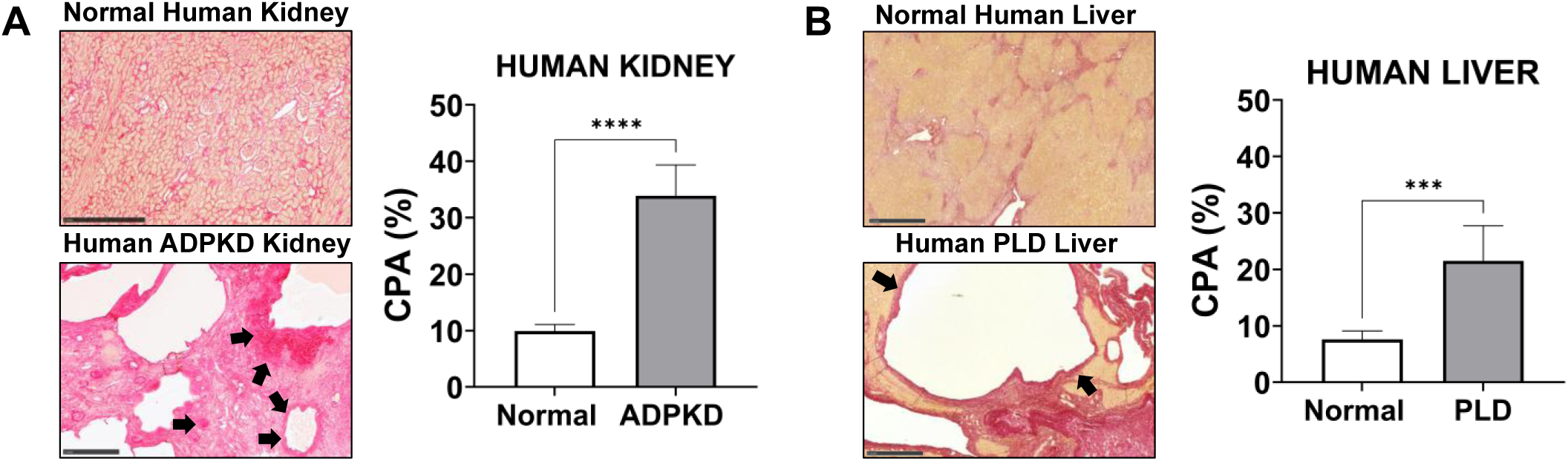
Collagen deposition is increased in human ADPKD kidney and polycystic liver disease tissue. **(A)** Representative Picrosirius red–stained sections of normal human kidney and autosomal dominant polycystic kidney disease (ADPKD) kidney, with quantification of collagen proportionate area (CPA). Arrows mark pericystic and interstitial collagen deposition. **(B)** Representative Picrosirius red–stained sections of normal human liver and polycystic liver disease (PLD) liver, with CPA quantification. Arrows mark collagen deposition surrounding hepatic cysts. Scale bars, 1 mm. CPA was quantified as the percentage of Sirius red–positive area in nonoverlapping fields per section at 4× magnification using ImageJ. Statistical significance was determined by two-tailed unpaired t-test with Welch’s correction. \*\*\**P* < 0.001, \*\*\*\**P* < 0.0001.

Consistent with previous reports (*27, 81, 82*), these data support collagen as a robust, disease-relevant molecular biomarker for both renal and hepatic fibrosis in ADPKD, providing a strong biological and translational rationale for collagen-targeted precision MRI approaches such as Gd-hProCA32.Collagen. Importantly, the spatial localization and quantitative elevation of collagen deposition underscore the importance of noninvasive detection, staging, and longitudinal monitoring of multi-organ fibrotic progression in ADPKD patients. Moreover, the multi-organ distribution of collagen overexpression uniquely positions Gd-hProCA32.Collagen as a pan-organ imaging agent capable of simultaneously detecting and staging fibrotic burden across the kidneys and liver in a single imaging session, an unmet capability in current ADPKD clinical practice.

### Development and Characterization of hProCA32.Collagen as a Collagen-Targeted Protein Contrast Agent for Precision MRI of Renal and Liver Fibrosis

To enable molecularly specific, noninvasive detection of hepatorenal fibrosis in ADPKD, we developed Gd-hProCA32.Collagen, an engineered protein-based MRI contrast agent incorporating two Gd³⁺ chelation sites and a high-affinity collagen-targeting moiety designed to selectively bind type I collagen enriched in fibrotic tissues (Fig. 3A). The protein scaffold is rationally engineered to achieve high longitudinal (r₁) and transverse (r₂) relaxivity through coordinated contributions from inner-sphere, second-sphere, and outer-sphere water interactions, optimized water exchange kinetics, and an increased rotational correlation time (τR) (*78–80, 83–85*).

**Fig. 3.**
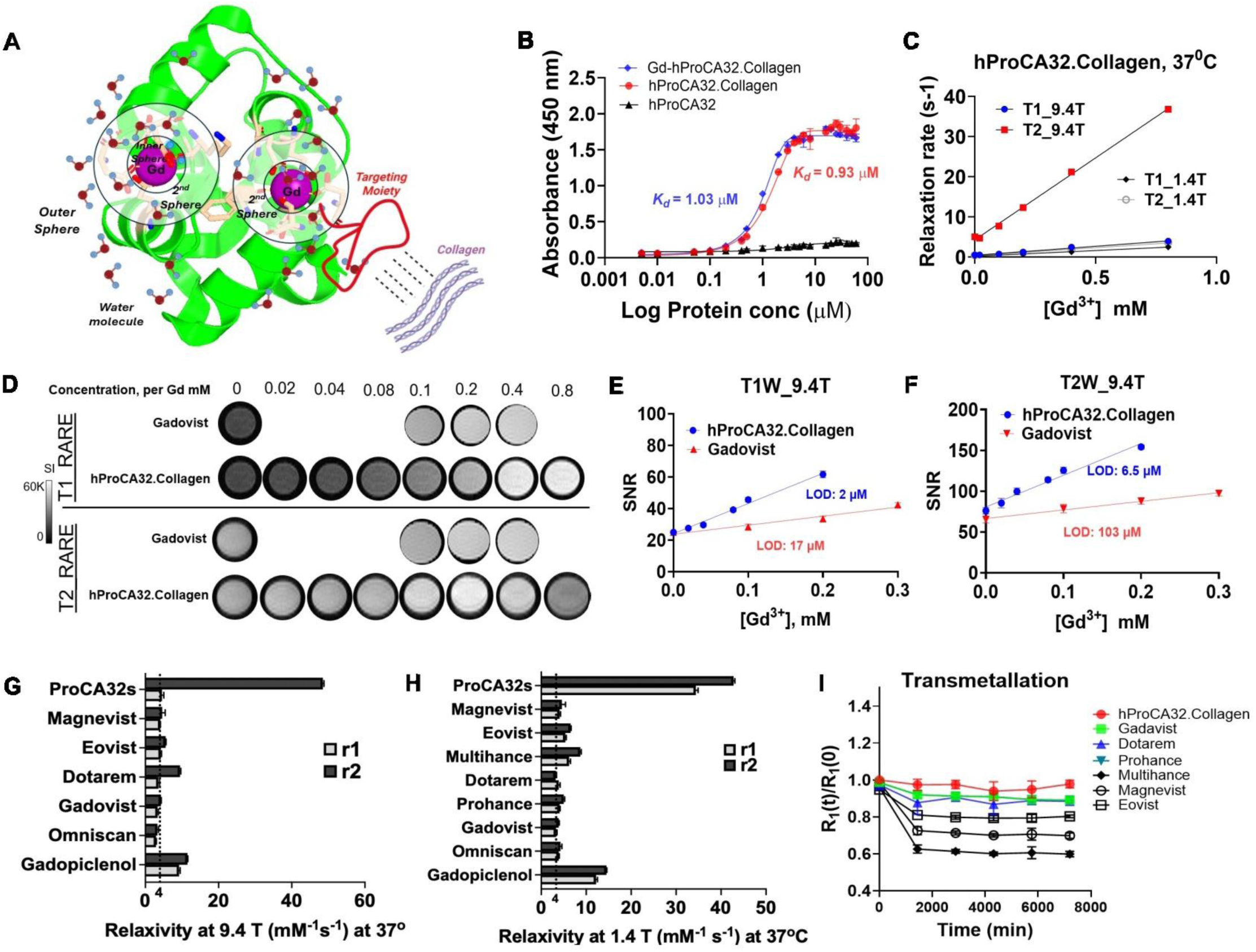
Design, collagen binding, relaxivity, and stability of Gd-hProCA32.Collagen. **(A)** Schematic of Gd-hProCA32.Collagen, showing the engineered protein scaffold with two Gd³⁺-binding sites, inner-, second-, and outer-sphere water molecules, and the collagen-targeting moiety bound to collagen fibrils. **(B)** Binding of Gd-hProCA32.Collagen, metal-free hProCA32.Collagen, and nontargeted hProCA32 to immobilized collagen type I, measured by enzyme-linked immunosorbent assay (ELISA) (absorbance at 450 nm). **(C)** Longitudinal (*R*₁) and transverse (*R*₂) relaxation rates of Gd-hProCA32.Collagen as a function of Gd concentration at 9.4 T and 1.4 T (37 °C). Relaxivities (r₁, r₂) were calculated from the slopes. **(D)** *T*₁-weighted and *T*₂-weighted rapid acquisition with relaxation enhancement (RARE) phantom images at 9.4 T of gadobutrol and Gd-hProCA32.Collagen at the indicated Gd concentrations. **(E and F)** Signal-to-noise ratio (SNR) versus Gd concentration on *T*₁-weighted (E) and *T*₂-weighted (F) images at 9.4 T. The limit of detection (LOD) was defined as the concentration producing an SNR increase of 3 SD above baseline. **(G and H)** Per-Gd r₁ and r₂ of Gd-hProCA32.Collagen compared with clinical gadolinium-based contrast agents (GBCAs) at 9.4 T (G) and 1.4 T (H), 37 °C. **(I)** Transmetallation stability, shown as the normalized longitudinal relaxation rate *R*₁(*t*)/*R*₁(0) of each agent over 7200 min in the presence of 2.5 mM ZnCl₂ in phosphate buffer (37 °C).

Binding studies demonstrate that both apo-and gadolinium (Gd)-loaded hProCA32.Collagen retained strong affinity for type I collagen, with dissociation constants of *K_d_* = 1.03 µM and 0.93 µM, respectively (Fig. 3B). In contrast, the non-targeted scaffold hProCA32 exhibited negligible binding across all tested concentrations (Fig. 3B), confirming the specificity conferred by the collagen-targeting domain. Binding affinity was preserved in rat, mouse, and dog serum and in human plasma (EC₅₀, 1.00 to 1.34 μM; fig. S1), supporting the translational potential of hProCA32.Collagen across preclinical species and into human clinical applications. At 1.4 T and 37 °C, Gd-hProCA32.Collagen had a per-Gd *r*₁ of 34.5 mM⁻¹ s⁻¹ and *r*₂ of 43.6 mM⁻¹ s⁻¹, about 3-fold higher than the highest-relaxivity clinical agent tested, gadopiclenol (Fig. 3, C and H). At 9.4 T, *r*₁ (4.53 mM⁻¹ s⁻¹) was similar to that of clinical GBCAs, whereas *r*₂ (48.2 mM⁻¹ s⁻¹) was about 5-to 10-fold higher (Fig. 3G). In phantoms at 9.4 T, the limits of detection were 2 μM for Gd-hProCA32.Collagen versus 17 μM for gadobutrol on *T*₁-weighted images and 6.5 μM versus 103 μM on *T*₂-weighted images, corresponding to 8.5-and 15.8-fold improvements in sensitivity (Fig. 3, D to F). Signal simulations were used to guide sequence selection for *in vivo* imaging at 9.4 T (fig. S2).

In a zinc transmetallation assay, Gd-hProCA32.Collagen retained 97% of its initial relaxation rate after 7200 min, comparable to macrocyclic agents (gadobutrol and gadoterate), but higher than linear agents (gadopentetate, gadobenate, and gadoxetate) (Fig. 3I). By SEC-HPLC, intact Gd-hProCA32.Collagen was detected in the serum of rats at 3 and 24 h after intravenous injection (0.010 mmol Gd/kg) but not at 48 or 72 h (fig. S3A). Serum [Gd] measured by ICP-OES showed the same pattern, peaking at 3 hours (139 ± 19.7 μM versus 38.4 ± 6.78 μM for gadobutrol at 0.1 mmol Gd/kg), then declining more slowly than for gadobutrol, consistent with the larger size of the protein agent, and returning to baseline by 72 h (fig. S3B). Gd-hProCA32.Collagen also remained predominantly monomeric, with no evidence of aggregation, after incubation in human serum at 37 °C for up to 14 days (fig. S3C).

Biodistribution measured by inductively coupled plasma optical emission spectroscopy (ICP-OES) showed that Gd accumulated mainly in the kidneys and liver, peaking at 24 h post-injection. Peak concentrations were higher in PKD rats than in control rats, consistent with the greater collagen content of cystic organs (fig. S4). By 72 h, Gd had largely cleared from control organs, whereas residual Gd remained in PKD liver (49.3 μM) and kidney (26.5 μM). Four days after injection, organ weights did not differ between Gd-hProCA32.Collagen–treated and saline-treated rats (*n* = 5 per group; *P* > 0.05), and no gross or histopathological abnormalities were observed on hematoxylin and eosin stained sections of the heart, liver, lung, spleen, or kidney (fig. S5).

Collectively, these results establish hProCA32.Collagen as a high-affinity, collagen-specific protein MRI contrast agent with substantially enhanced relaxivity, sensitivity, and stability compared to conventional GBCAs, providing a strong mechanistic and biophysical foundation for its use in precision MRI (pMRI) of renal fibrosis in ADPKD.

### Conventional clinical indices do not detect early fibrosis in PCK rats

The PCK rat, which carries a spontaneous *Pkhd1* mutation, develops progressive renal and hepatic cysts and has been widely used in preclinical studies of ADPKD therapeutics, including the studies that preceded clinical testing of tolvaptan (*48, 86, 87*). This model shows conserved ciliary localization defects, dysregulated downstream signaling pathways, and strong phenotypic concordance with human disease, and has served as a translational platform in preclinical studies supporting therapies later evaluated in clinical trials (*88, 89*). To enable comparison with genetically defined controls, PCK rats were backcrossed onto the Charles River International Genetic Standardization (IGS) Sprague-Dawley (SD) background, and WT IGS SD rats were used as controls. Histopathological analysis with picrosirius red staining revealed pericystic collagen deposition in both kidney and liver of PCK rats at 26 weeks (Fig. 4A), recapitulating the combined renal and hepatic involvement in human polycystic disease.

**Fig. 4.**
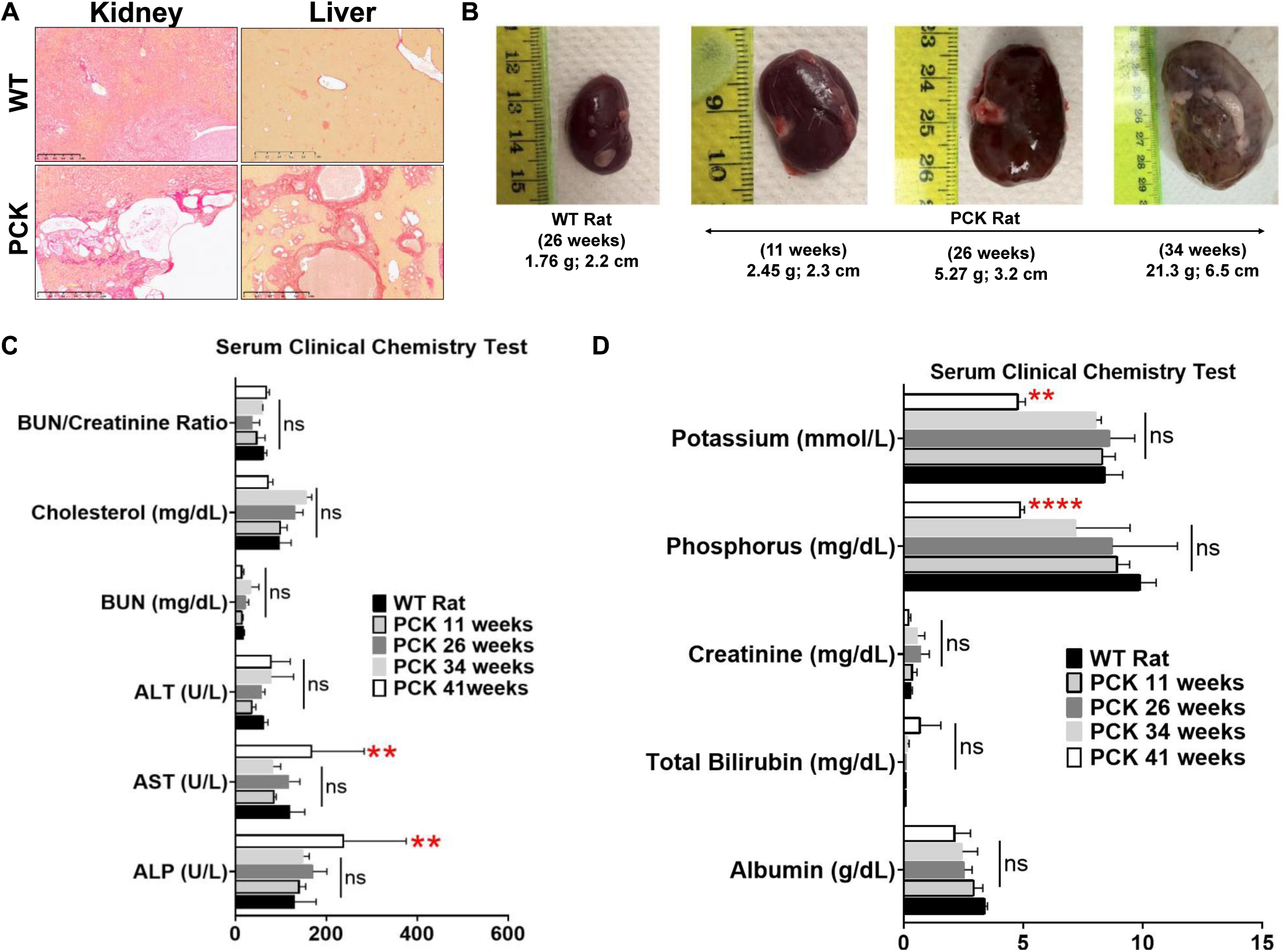
Cystic disease progresses in PCK rats while serum markers of kidney and liver function remain normal until late stages. **(A)** Representative Picrosirius red–stained sections of kidney and liver from WT and PCK rats at 26 weeks of age, showing cysts surrounded by collagen deposition in PCK rats (Scale bars, 1 mm). **(B)** Representative gross images of kidneys from a WT rat at 26 weeks and PCK rats at 11, 26, and 34 weeks of age. Values below each image are the weight and length of the kidney shown. (**C and D**) Serum clinical chemistry in WT rats (26 weeks) and PCK rats at 11, 26, 34, and 41 weeks of age. (C) BUN/creatinine ratio, cholesterol, blood urea nitrogen (BUN), alanine aminotransferase (ALT), aspartate aminotransferase (AST), and alkaline phosphatase (ALP). (D) Potassium, phosphorus, creatinine, total bilirubin, and albumin. Data are means ± SD; n = 7 rats per group. Statistical significance versus WT was determined by one-way ANOVA with Dunnett’s multiple comparisons test. \*\**P* < 0.01, \*\*\*\**P* < 0.0001; ns, not significant.

To determine whether conventional measures reflect this fibrotic remodeling, kidney morphology and serum chemistry were assessed in PCK rats at 11, 26, 34, and 41 weeks of age and in WT rats at 26 weeks (*n* = 7 per group; Fig. 4, B to D, and table S1). Kidneys enlarged progressively with age in PCK rats, with kidney weight increasing from 2.45 g at 11 weeks to 21.3 g at 34 weeks in representative animals (Fig. 4B). Despite this, serum markers of glomerular filtration did not distinguish PCK from WT rats at any age. Serum creatinine (0.23 to 0.71 mg/dl versus 0.30 mg/dl in WT), blood urea nitrogen (BUN; 16.0 to 35.0 versus 18.7 mg/dl), and BUN/creatinine ratio did not differ from WT at any time point (Fig. 4, C and D, and table S1). This is consistent with the preserved renal function that characterizes the early and intermediate stages of human ADPKD despite ongoing cyst growth.

Serum liver chemistry followed a similar pattern. ALT, total bilirubin, albumin, and cholesterol did not differ from WT at any age, whereas ALP and AST were increased only at 41 weeks (ALP, 240 versus 131 U/liter; AST, 169 versus 121 U/liter; *P* < 0.01) (Fig. 4C). Because ALT and bilirubin remained normal, the increase in ALP is consistent with biliary involvement associated with progressive hepatic cystic disease rather than with hepatocellular injury, whereas the isolated increase in AST is nonspecific. Serum potassium and phosphorus differed from WT only at 41 weeks (potassium, 4.8 versus 8.4 mmol/liter, *P* < 0.01; phosphorus, 4.9 versus 9.9 mg/dl, *P* < 0.0001) (Fig. 4D). Other serum parameters, including electrolytes, proteins, lipids, and pancreatic enzymes, exhibited no consistent disease-related pattern (table S1).

Thus, in PCK rats, pericystic fibrosis is established while serum markers of kidney and liver function remain unchanged, and these markers change only at advanced disease stages. Conventional serum chemistry therefore does not capture fibrotic remodeling during the period when intervention would be most beneficial.

### PCK Rats Exhibit Progressive Renal Enlargement and Increased Kidney-to-Body Weight Ratio Despite Comparable Body and Liver Weight to Wild-Type Controls

Consistent with previous reports (*90, 91*), PCK rats exhibited progressive kidney and liver enlargement (Fig. 5). At 11 weeks, kidney weight and KW/BW were modestly increased in PCK rats compared with WT rats (*P* < 0.05). This partly reflects the lower body weight of PCK rats at this age (*P* < 0.01) (Fig. 5, A, B, and D). Liver weight, LW/BW, and TKV/BW did not differ from WT at 11 weeks (Fig. 5, C, E, and F). By 26 weeks, body weight no longer differed between groups (*P* = 0.83) (Fig. 5A), whereas kidney weight, KW/BW, liver weight, LW/BW, and TKV/BW were all increased in PCK rats (*P* < 0.01) (Fig. 5, B to F). TKV normalized to body length did not differ at either age (*P* = 0.41 at 11 weeks and *P* = 0.18 at 26 weeks) (Fig. 5G). MRI-derived TKV followed the same pattern. TKV measured on *T*₂-weighted images did not differ from WT at 11 weeks (*P* = 0.17) but was increased at 26 weeks (*P* = 0.014) (Fig. 5H). TKV measured on *T*₁-weighted images gave concordant results (*P* < 0.05 at 26 weeks) (Fig. 5I).

**Fig. 5.**
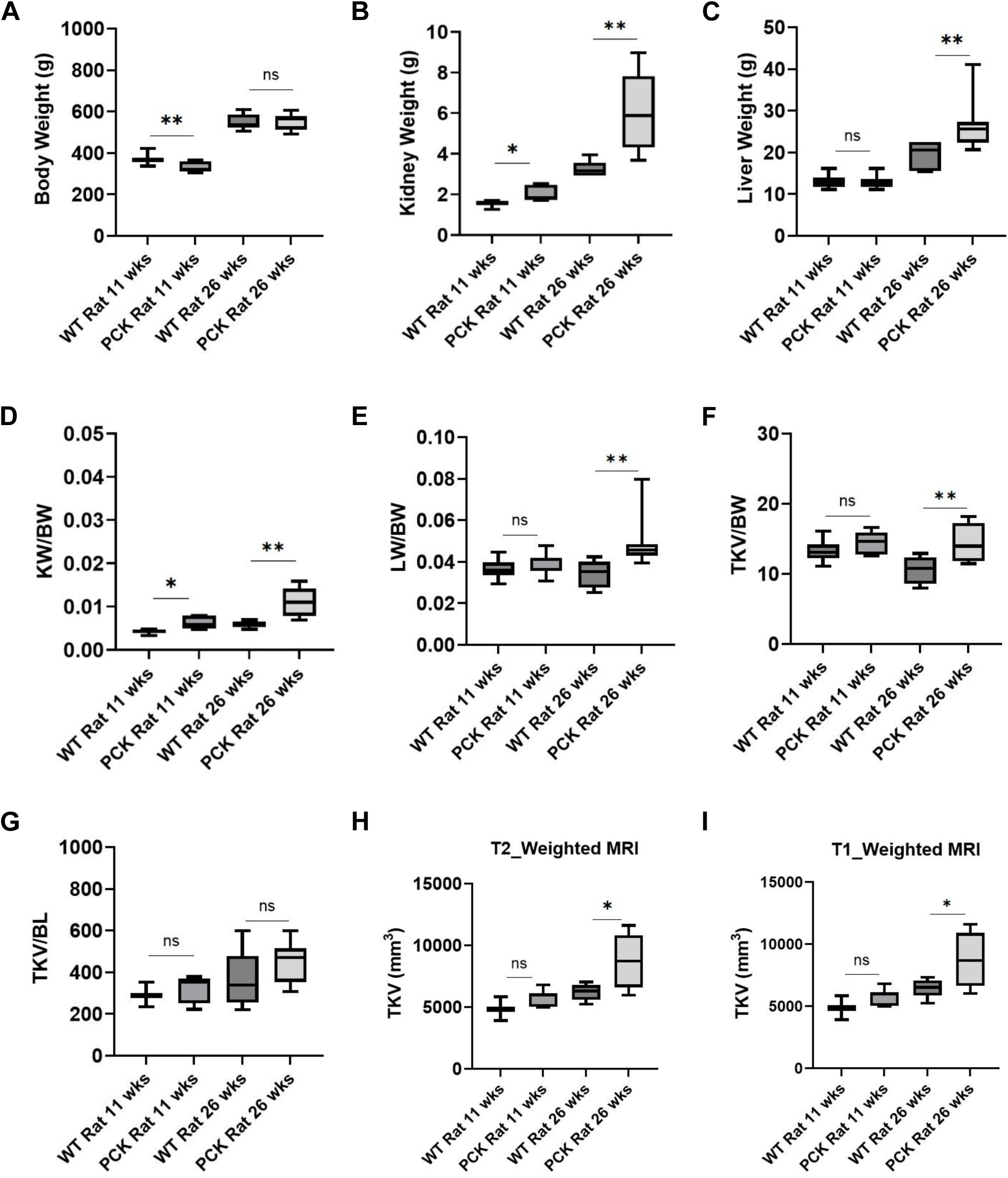
Morphometric and MRI-derived volumetric analysis of WT and PCK rats at early (11 weeks) and moderately advanced disease (26 weeks). (**A to C**) Body weight (A), single-kidney weight (B), and liver weight (C). (**D and E**) Kidney weight–to–body weight ratio (KW/BW) (D) and liver weight–to–body weight ratio (LW/BW) (E). (**F and G**) Total kidney volume (TKV) normalized to body weight (TKV/BW) (F) and to body length (TKV/BL) (G). (**H and I**) TKV measured on *T*₂-weighted (H) and *T*₁-weighted (I) MRI. Box plots show the median and interquartile range; whiskers show minimum and maximum. *n* = 5 WT and 8 PCK rats at 11 weeks and 5 WT and 10 PCK rats at 26 weeks. PCK and age-matched WT rats were compared by two-tailed Welch’s *t-test*. \**P* < 0.05, \*\**P* < 0.01; ns, not significant.

Importantly, while MRI-derived TKV detected kidney enlargement at 26 weeks, it provided no information on the tissue processes that drive disease progression, such as fibrotic remodeling and pericystic collagen deposition, which may already be present at 11 weeks, when TKV does not yet differ from WT. Gross anatomical measures such as TKV, therefore, cannot detect the early fibrotic changes that precede organ enlargement. This limitation motivates molecular imaging approaches that directly quantify collagen deposition *in vivo*, such as the collagen-targeted pMRI strategy evaluated in this study.

### Collagen-targeted MRI provides sustained renal enhancement and improves cyst detection in PCK rats

We next evaluated Gd-hProCA32.Collagen *in vivo* in 26-week-old PCK and WT rats imaged before and after intravenous injection of Gd-hProCA32.Collagen at 0.010 mmol Gd/kg or the clinical agent gadobutrol (Gadovist) at 0.1 mmol Gd/kg, a 10-fold higher gadolinium dose (fig. S6). PCK rats receiving Gd-hProCA32.Collagen (*n* = 15) or Gadovist (*n* = 9) did not differ in baseline cyst count, total cyst volume, total kidney volume (TKV), or cyst index (fig. S7 and table S2).

In PCK kidneys, Gd-hProCA32.Collagen increased SNR progressively, reaching 39% on *T*₁-weighted and 49% on *T*₂-weighted images 48 h after injection. Gadovist produced a small, transient increase (≤10%) that peaked within 3 h and returned toward baseline by 24 h (Fig. 6, A and B, and fig. S8A). AUC Mean enhancement over 48 h was higher with Gd-hProCA32.Collagen than with Gadovist on both *T*₁-weighted (26.3 ± 1.4% versus 5.1 ± 0.2 %; *P* < 0.01) and *T*₂-weighted images (41.6 ± 0.8% versus 4.2 ± 0.1%; *P* < 0.001) (Fig. 6D). Transverse relaxation rate measurements were consistent with tissue retention. With Gd-hProCA32.Collagen, ΔR₂ in cysts increased from 5.6 s⁻¹ at 30 min to 13.8 s⁻¹ at 48 h, and noncystic parenchyma exhibited a similar increase (3.4 to 10.1 s⁻¹). With Gadovist, ΔR₂ declined to near baseline by 24 h (fig. S9).

**Fig. 6.**
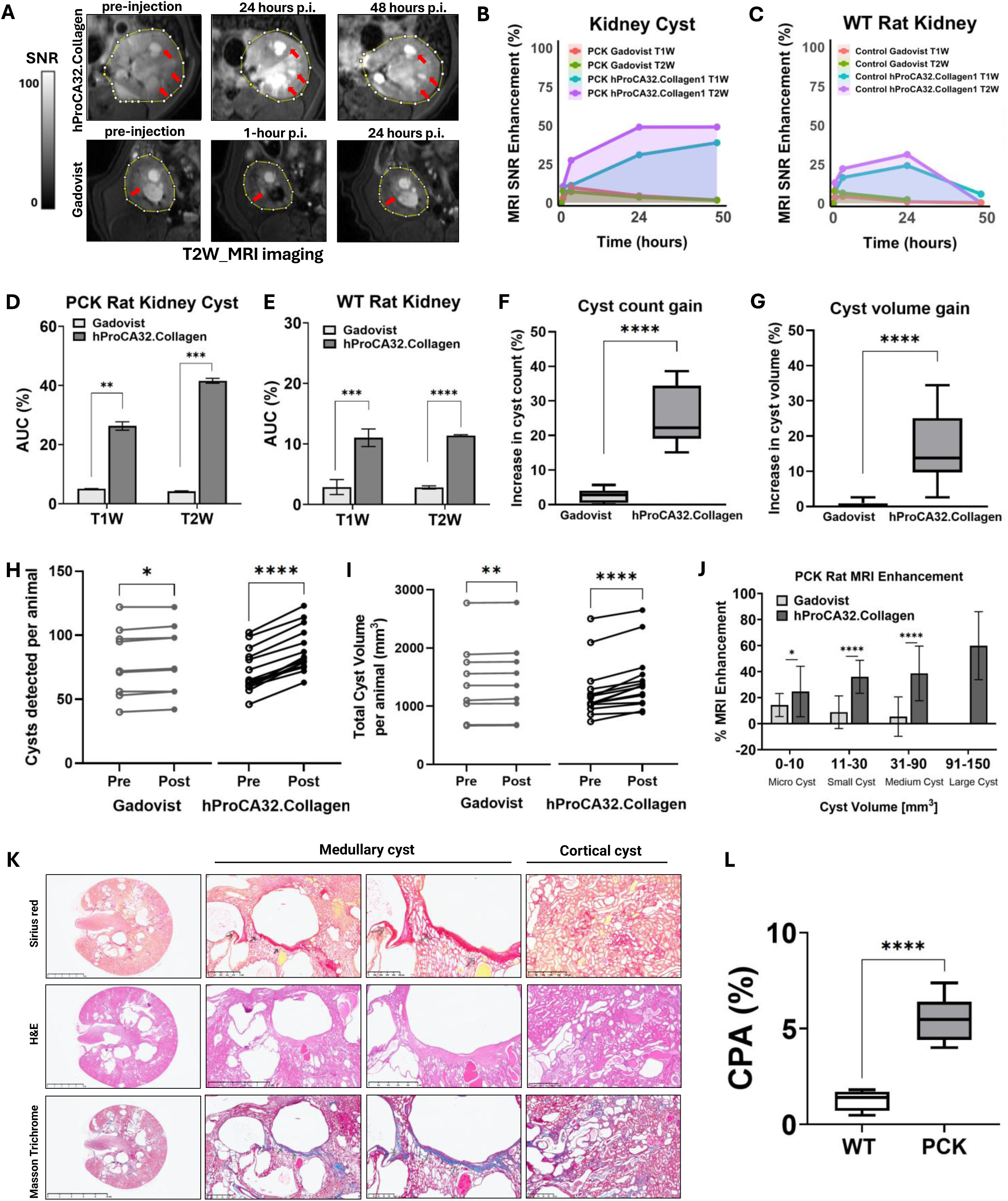
Collagen-targeted MRI provides sustained renal enhancement and improves cyst detection in PCK rats. (**A**) Representative *T*₂-weighted MRI signal-to-noise ratio (SNR) maps of PCK rat kidneys (yellow outlines) before and 24 and 48 h after injection of Gd-hProCA32.Collagen (top), and before and 1 and 24 h after injection of Gadovist (gadobutrol; bottom). Arrows indicate enhancing cysts. (**B and C**) Percent SNR enhancement over time on T₁-and T₂-weighted images in PCK rat kidney cysts (B) and WT rat kidneys (C). (**D and E**) Area under the enhancement–time curve (AUC) in PCK rat kidney cysts (D) and WT rat kidneys (E). AUC was calculated by the trapezoidal rule from 0 to 48 h and is expressed as mean enhancement over 48 h (AUC/48 h, %). (**F and G**) Percent increase in detected cyst count (F) and total cyst volume (G) in each PCK rat after contrast. (**H and I**) Number of detected cysts (H) and total cyst volume (I) in each PCK rat before (open circles) and after (filled circles) contrast; lines connect values from the same rat. (**J**) Percent signal enhancement by cyst size category in PCK rats; no large cysts (91 to 150 mm³) were detected after Gadovist. (**K**) Representative Picrosirius red, hematoxylin and eosin (H&E), and Masson’s trichrome stained sections of a PCK rat kidney showing collagen deposition surrounding medullary and cortical cysts. Scale bars, 5 mm (left column), 1 mm (second column), and 500 µm (third and fourth columns). (**L**) Collagen proportionate area (CPA) in WT and PCK rat kidneys. All rats were 26 weeks old. Gd-hProCA32.Collagen was given at 0.010 mmol Gd/kg and Gadovist at 0.1 mmol Gd/kg; cysts were quantified 24 h (Gd-hProCA32.Collagen) or 1 h (Gadovist) after injection. *n* = 15 (Gd-hProCA32.Collagen) and 9 (Gadovist) PCK rats and 5 WT rats per agent; *n* = 5 WT and 10 PCK rats in (L). Data are mean ± SD (B to E and J) or box plots showing median, interquartile range, and minimum to maximum (F, G, and L). Comparisons used the Mann–Whitney test (C, E to G, J, and L) and the Wilcoxon matched-pairs signed rank test (H and I). \**P* < 0.05, \*\**P* < 0.01, \*\*\**P* < 0.001, \*\*\*\**P* < 0.0001, post-injection.

Three observations indicated that sustained enhancement reflected collagen binding rather than slow clearance alone. First, in WT kidneys (*n* = 5 per agent), Gd-hProCA32.Collagen enhancement peaked at 24 h (24.2% on *T*₁-weighted and 31.2% on *T*₂-weighted images) and returned to baseline by 48 h (Fig. 6C and fig. S10). AUC Mean enhancement in WT kidneys (11.3% and 10.5%) was about one-third of that in PCK kidneys, although it still exceeded that of Gadovist (*P* < 0.001 and *P* < 0.0001) (Fig. 6E). Second, the nontargeted protein agent, hProCA32, given at the same gadolinium dose, also enhanced PCK kidneys but peaked at 24 h and declined by 48 h (retention index, 0.77 on *T*₁-weighted and 0.64 on *T*₂-weighted images, versus 1.00 for Gd-hProCA32.Collagen). AUC Mean enhancement over 48 h was therefore 2.0-to 2.3-fold higher with the targeted agent. Similar results were obtained in the liver (retention index, 0.52 to 0.53 versus 0.88 to 0.95; 1.8-fold higher mean enhancement) (fig. S11 and table S3). Third, PCK kidneys exhibited dense collagen surrounding medullary and cortical cysts, and CPA was higher than in WT kidneys (5.5 ± 1.2% versus 1.3 ± 0.6%; *P* < 0.0001) (Fig. 6, K and L).

Because Gd-hProCA32.Collagen enhances collagen-rich pericystic matrix, the next question to ask was whether it improves cyst delineation. Cyst count and total cyst volume increased after both agents: *P* < 0.0001 for both measures with Gd-hProCA32.Collagen, and *P* < 0.05 (count) and *P* < 0.01 (volume) with Gadovist (Fig. 6, H and I). The gain was much larger with Gd-hProCA32.Collagen, however, which revealed 17.0 ± 3.9 additional cysts per animal compared with 1.8 ± 1.2 for Gadovist. This corresponds to increases in detected cyst number of 25.1 ± 8.4% versus 2.6 ± 2.0% (*P* < 0.0001) and in delineated cyst volume of 16.3 ± 10.1% versus 0.7 ± 0.8% (*P* < 0.0001) (Fig. 6, F and G, and table S2). The detection gain ranged from 15 to 39% in rats receiving Gd-hProCA32.Collagen and from 0 to 6% in rats receiving Gadovist, with no overlap between groups. Enhancement was greater with Gd-hProCA32.Collagen for microcysts (0 to 10 mm³; 24.7% versus 14.3%; *P* < 0.05), small cysts (11 to 30 mm³; 36.0% versus 8.8%; *P* < 0.0001), and medium cysts (31 to 90 mm³; 38.6% versus 5.4%; *P* < 0.0001). Large cysts (91 to 150 mm³) were delineated only with Gd-hProCA32.Collagen (60% enhancement) (Fig. 6J). Across individual cysts, enhancement with Gd-hProCA32.Collagen was moderately associated with cyst volume (*r* = 0.37), whereas Gadovist enhancement was not (*r* = 0.09) (fig. S8B).

Finally, *T*₂-weighted enhancement 24 h after Gd-hProCA32.Collagen completely separated PCK (*n* = 15) from WT (*n* = 5) kidneys (area under the ROC curve, 1.00; 95% CI, 1.00 to 1.00; *P* = 0.007). Gadovist enhancement at its peak did not significantly discriminate PCK (*n* = 9) from WT (*n* = 5) kidneys (0.93; 95% CI, 0.75 to 1.00; *P* = 0.053) (fig. S12).

### Collagen-targeted MRI maps striation-like fibrosis associated with microcyst clusters

In addition to pericystic enhancement, Gd-hProCA32.Collagen revealed radially oriented, striation-like enhancement extending from the medulla toward the renal capsule in PCK kidneys, which was not seen with Gadovist (Fig. 7A). *Ex vivo* MRI of excised kidneys showed the same radial pattern at higher resolution (fig. S13A). In these regions, enhancement peaked at 24 h (33.0% on *T*₁-weighted and 37.1% on *T*₂-weighted images) and remained elevated at 48 h (27.4% and 36.1%), whereas Gadovist produced less than 8% enhancement at all time points (Fig. 7B). AUC Mean enhancement over 48 h was about 11-fold higher with Gd-hProCA32.Collagen than with Gadovist (25.1 ± 0.7% versus 2.4 ± 0.4% on *T*₁-weighted images, *P* < 0.0001; 29.3 ± 0.3% versus 2.8 ± 0.2% on *T*₂-weighted images, *P* < 0.001; *n* = 15 and 9) (Fig. 7C).

**Fig. 7.**
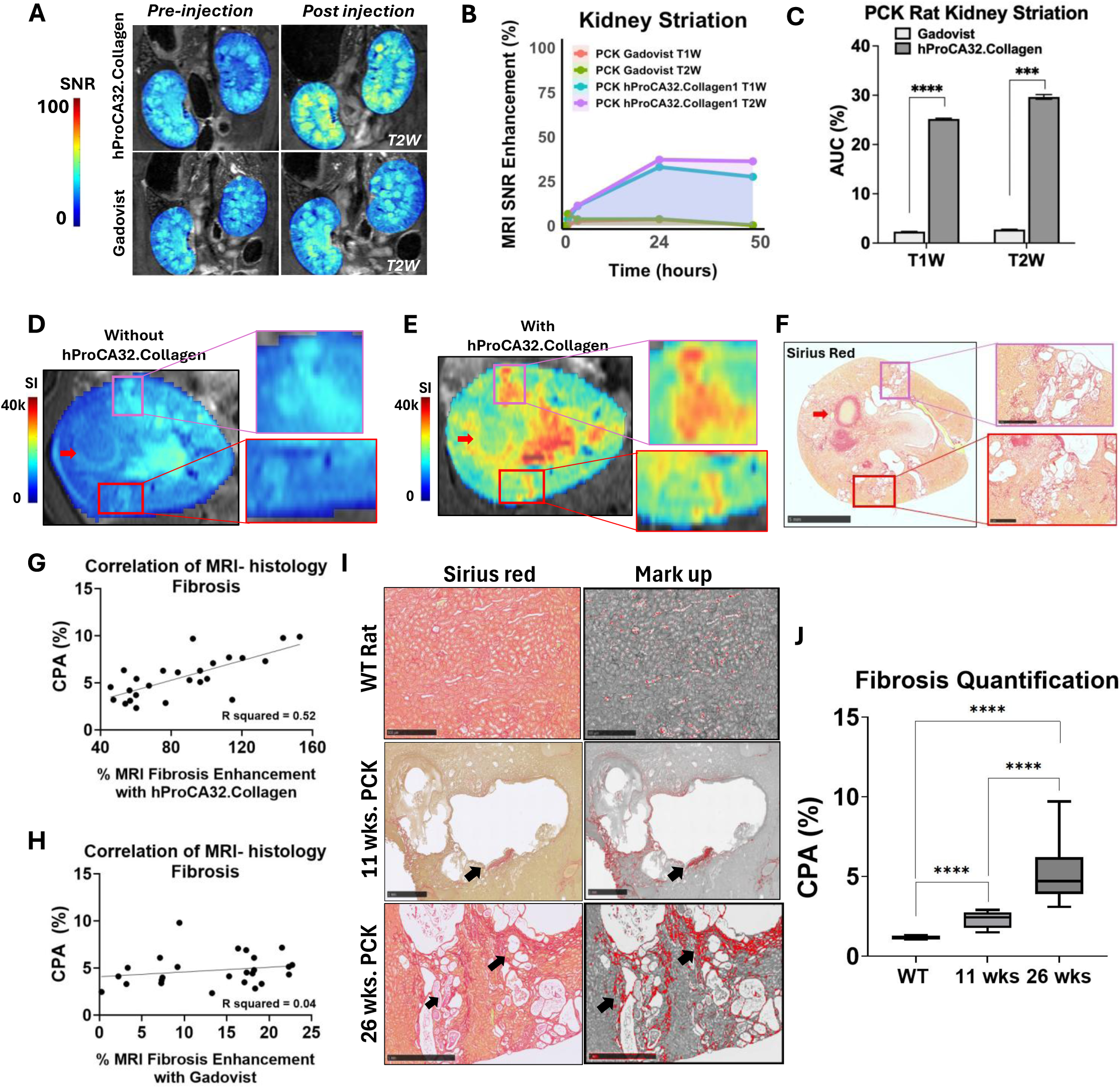
Collagen-targeted MRI detects pericystic and cortical fibrosis in PCK rats, with histological validation. (**A**) Representative *T*₂-weighted SNR maps of PCK rat kidneys before and 24 h after Gd-hProCA32.Collagen (top) or before and 1 h after Gadovist (bottom), showing radially oriented, striation-like enhancement after Gd-hProCA32.Collagen. (**B and C**) Percent enhancement over time on *T*₁-and *T*₂-weighted images (B) and area under the enhancement–time curve (AUC) (C) in striation regions of PCK kidneys; AUC is expressed as mean enhancement over 48 h (AUC/48 h, %). **(D to F**) Co-registered *T*₂-weighted signal intensity (SI) maps of the same PCK kidney before (D) and 24 h after (E) Gd-hProCA32.Collagen, and the corresponding Picrosirius red–stained section (F). Boxes indicate magnified regions; red arrows mark corresponding landmarks. (**G and H**) Correlation between regional enhancement and collagen proportionate area (CPA) after Gd-hProCA32.Collagen (G) or Gadovist (H). Each point represents one cortical or OSOM region of interest matched to the corresponding region on Picrosirius red–stained sections; regional enhancement values therefore exceed whole-kidney averages. Lines show linear regression. (**I**) Picrosirius red–stained sections (left) and collagen segmentation masks (right) from WT rats and PCK rats at 11 and 26 weeks; arrows indicate pericystic collagen. (**J**) Renal CPA in WT rats (26 weeks) and PCK rats at 11 and 26 weeks. Scale bars are shown. *n* = 15 (Gd-hProCA32.Collagen) and 9 (Gadovist) PCK rats in (B) and (C); 25 regions from 5 rats per agent in (G) and (H); 5 rats per group in (J). Data are mean ± SD (B and C) or box plots showing median, interquartile range, and minimum to maximum (J). Comparisons used the Mann–Whitney test (C), two-tailed Pearson correlation (G and H), and Kruskal–Wallis with Dunn’s multiple comparisons test (J). \*\*\**P* < 0.001, \*\*\*\**P* < 0.0001.

Co-registration of *in vivo* MRI with histology from the same kidney showed regions of focal enhancement after administration of Gd-hProCA32.Collagen, which corresponded to clusters of microcysts surrounded by Picrosirius red–positive collagen (Fig. 7, D to F). Correspondence was regional rather than pixel-level, reflecting differences in resolution and section plane between MRI and histology. These striations lay mainly in the cortex, where cysts were smaller than in the outer stripe of the outer medulla (OSOM) (8.2 ± 10.7 versus 16.8 ± 17.6 mm³) yet fibrosis was greater. The cortex showed a higher fibrosis index measured on whole regions (0.13 ± 0.02 versus 0.02 ± 0.03; *P* < 0.0001) and a higher CPA measured in selected fields (3.7 ± 1.5% versus 2.2 ± 0.9%; *P*< 0.05; *n* = 10 rats) than the OSOM (fig. S13, B to D). WT kidneys showed normal architecture with minimal collagen (fig. S13, E and F).

Across cortical and OSOM regions, enhancement after Gd-hProCA32.Collagen correlated with CPA measured in matched histological regions (r = 0.72; 95% CI, 0.46 to 0.87; P < 0.0001; n = 25 regions from 5 rats). In contrast, enhancement after Gadovist was small (0 to 22%) and did not correlate significantly with CPA (r = 0.21; 95% CI, −0.21 to 0.56; P = 0.33; n = 25 regions from 5 rats) (Fig. 7, G and H). Finally, we examined when fibrosis develops relative to kidney enlargement. Collagen deposition around cysts was already evident in 11-week-old PCK rats and became extensive by 26 weeks (Fig. 7I). Consistent with the possibility raised above, renal CPA was already increased at 11 weeks (2.3 ± 0.5% versus 1.2 ± 0.1% in WT rats; P < 0.0001), when TKV did not yet differ from WT (Fig. 5, H and I). CPA rose further by 26 weeks (5.2 ± 1.7%; P < 0.0001 versus 11 weeks; n = 10 rats per group) (Fig. 7J). Thus, pericystic fibrosis precedes measurable kidney enlargement in PCK rats, and collagen-targeted MRI detected and mapped this fibrotic component, which volumetric imaging does not capture.

### Collagen-targeted MRI detects hepatic fibrosis in PCK rats

Polycystic liver disease with pericystic fibrosis frequently accompanies ADPKD (Fig. 2B), so it was important to determine whether Gd-hProCA32.Collagen could assess the liver in the same imaging session. In PCK rat livers, Gd-hProCA32.Collagen produced enhancement that rose to 51.0% on *T*₁-weighted and 64.3% on *T*₂-weighted images at 24 h and remained elevated at 48 h (49.0% and 56.4%). Gadovist enhancement was less than 5.8% and returned to baseline by 24 h (Fig. 8, A to C). AUC Mean enhancement over 48 h was higher with Gd-hProCA32.Collagen than with Gadovist (25 ± 2.3% versus 1.7 ± 0.3% on *T*₁-weighted images and 36.3 ± 1.2% versus 2.6 ± 0.4% on *T*₂-weighted images; both *P* < 0.0001; *n* = 15 and 9) (Fig. 8D).

**Fig. 8.**
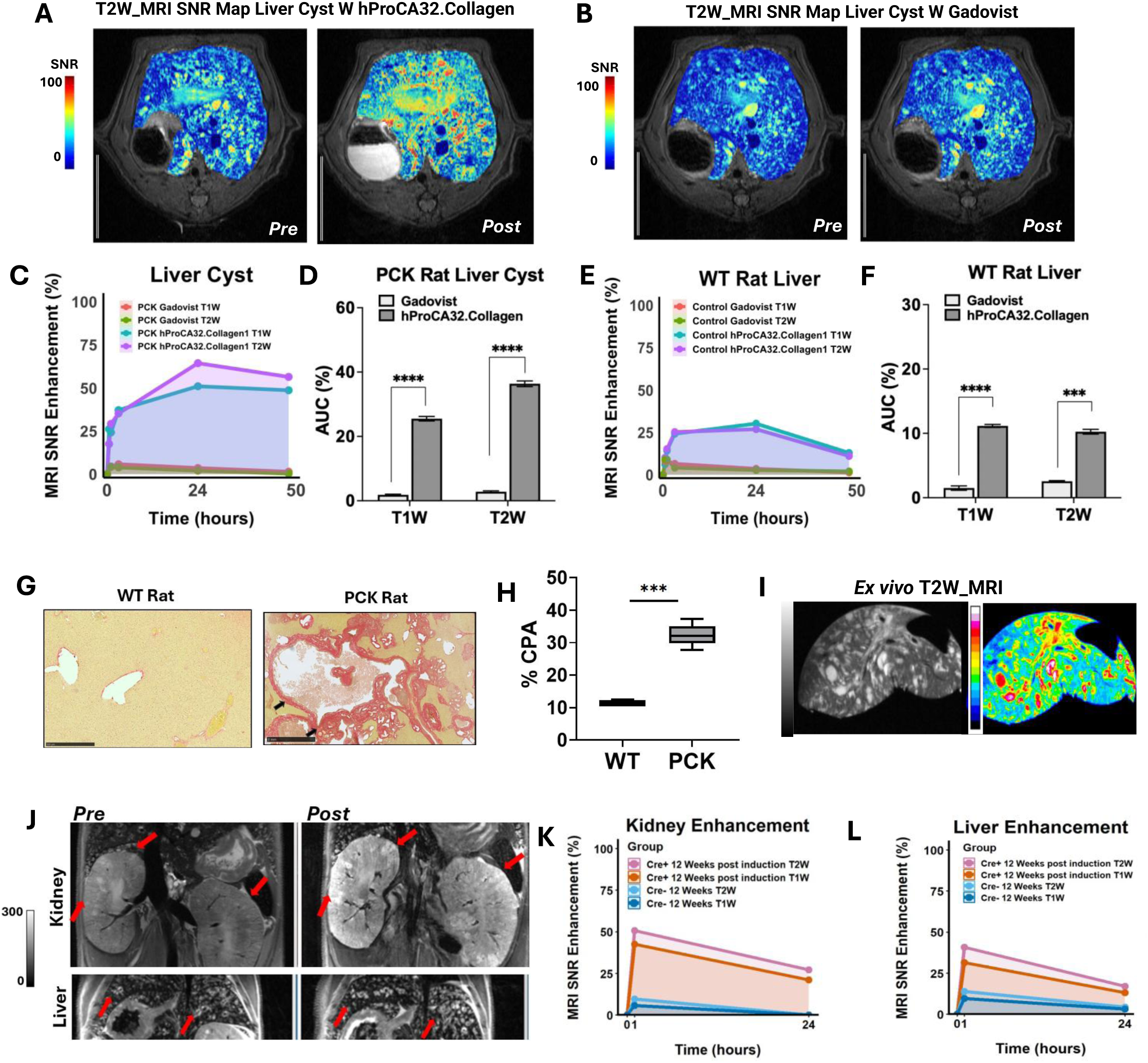
Collagen-targeted MRI detects hepatic fibrosis in PCK rats and renal and hepatic fibrosis in *Pkd2* mutant mice. (**A and B**) Representative T₂-weighted SNR maps of PCK rat livers before and 24 h after Gd-hProCA32.Collagen (A) or before and 1 h after Gadovist (B). (**C and D**) Percent enhancement over time on T₁-and T₂-weighted images (C) and AUC (D) in PCK rat livers. (**E and F**) Corresponding enhancement time course (E) and AUC (F) in WT rat livers. AUC is expressed as mean enhancement over 48 h (AUC/48 h, %). (**G**) Representative Picrosirius red–stained liver sections from WT and PCK rats; arrows indicate pericystic collagen. (**H**) Hepatic collagen proportionate area (CPA) in WT and PCK rats. (**I**) *Ex vivo* T₂-weighted image (top) and SNR map (bottom) of an excised PCK rat liver. (**J**) Representative T₂-weighted images of the kidney (top) and liver (bottom) of a Cre⁺ *Pkd2* mutant mouse 12 weeks after tamoxifen induction, before and 1 h after Gd-hProCA32.Collagen (0.010 mmol Gd/kg); arrows indicate enhancing cysts. (**K and L**) Percent enhancement in the kidney (K) and liver (L) of Cre⁺ and Cre⁻ mice on T₁-and T₂-weighted images 1 and 24 h after injection. Rats were 26 weeks old; Gd-hProCA32.Collagen was administered at 0.010 mmol Gd/kg and Gadovist at 0.1 mmol Gd/kg. n = 15 (Gd-hProCA32.Collagen) and 9 (Gadovist) PCK rats and 5 WT rats per agent in (C) to (F); 5 rats per group in (H); 2 Cre⁺ and 2 Cre⁻ mice in (K) and (L). Data are mean ± SD (C to F, K, and L) or box plots showing median, interquartile range, and minimum to maximum (H). Mann–Whitney test. ***P < 0.001, ****P < 0.0001. Scale bars are shown.

In WT livers (*n* = 5 per agent), Gd-hProCA32.Collagen enhancement peaked at 24 h and declined by 48 h. AUC Mean enhancement (11.3 ± 0.3% and 10.5 ± 0.5%) was about one-third of that in PCK livers, although it still exceeded that of Gadovist (*P* < 0.0001 and *P* < 0.001) (Fig. 8, E and F). Consistent with collagen-dependent retention, the nontargeted protein hProCA32 retained only about half of its peak hepatic enhancement at 48 h, compared with 88 to 95% for Gd-hProCA32.Collagen (fig. S11B and table S3).

Picrosirius red staining showed dense collagen surrounding hepatic cysts in PCK rats, and hepatic CPA was about threefold higher than in WT rats (32.4 ± 3.4% versus 11.3 ± 1.0%; *n* = 5 per group; *P* < 0.001) (Fig. 8, G and H). This parallels the increase in CPA observed in human polycystic liver (Fig. 2B). *Ex vivo* MRI of the excised PCK liver revealed heterogeneous signal consistent with the distribution of cysts and fibrotic tissue (Fig. 8I). Serum markers of liver function did not differ from WT through 34 weeks of age (Fig. 4, C and D, and table S1), even though hepatic fibrosis was already established. Thus, collagen-targeted MRI detected hepatic fibrosis during a phase when conventional serum markers are uninformative, allowing the kidney and liver to be assessed in a single examination.

### Collagen-targeted MRI detects renal and hepatic fibrosis in an orthologous *Pkd2* mouse model

To test whether these findings extend to a model carrying an ADPKD gene mutation, a proof-of-concept study in tamoxifen-inducible *Pkd2* knockout mice (*92, 93*) was performed. Male mice were induced at 4 weeks of age, and uninduced Cre⁻ littermates served as controls. In a longitudinal cohort, Cre⁺ mice developed progressive kidney enlargement. TKV increased from 518.3 mm³ at 4 weeks after induction to about 3,805 mm³ at 20 weeks, compared with about 450 mm³ in Cre⁻ mice, and kidney weight at 20 weeks was about 10-fold higher. Body weight was similar until 20 weeks, when it was lower in Cre⁺ mice (fig. S14, A to C; *n* = 2 per group).

Twelve weeks after induction, Gd-hProCA32.Collagen (0.010 mmol Gd/kg) increased kidney signal by 44.6% on *T*₁-weighted and 50.7% on *T*₂-weighted images and liver signal by 32.0% and 40.7% at 1 h after injection. Enhancement remained above that of Cre⁻ controls at 24 h, whereas enhancement in Cre⁻ mice was 5 to 13% (Fig. 8, J to L, and fig. S14D; *n* = 2 per group). Enhancement peaked earlier in mice than in rats, consistent with faster clearance in the smaller animal. Kidneys and livers collected after the 24-h scan exhibited pericystic collagen on Picrosirius red staining, with higher CPA in Cre⁺ than in Cre⁻ mice in both organs (kidney, 1.3 versus 0.65%; liver, 0.8 versus 0.3%) (fig. S14, E and F). Although a smaller number of mice were involved in the proof-of-concept study, the results were consistent with the findings in PCK rats.

## Discussion

Results of this study demonstrated the *in vivo* application of a collagen-targeted, protein-based precision MRI platform for simultaneous molecular detection, enabling spatial mapping and staging of renal and hepatic fibrosis in rodent models of polycystic kidney disease. Gd-hProCA32.Collagen enabled visualization of subclinical extracellular matrix remodeling, pathological collagen deposition, and cyst-associated cortical fibrosis in very early disease stages when conventional functional biomarkers and approved GBCA remained indiscriminate. In PCK rats, pericystic collagen deposition was already increased at 11 weeks of age, before MRI-measured TKV changed and long before serum markers of kidney and liver function were altered. At 26 weeks, Gd-hProCA32.Collagen produced sustained kidney and liver enhancement at one-tenth the gadolinium dose of Gadovist, improved cyst delineation, revealed striation-like enhancement at microcyst clusters, and generated signal that correlated with histological collagen. Together, these findings indicate that collagen-targeted MRI can detect a fibrotic component of cystic disease that volumetric imaging and serum chemistry do not capture.

The results also indicated that the sustained enhancement reflects collagen binding rather than slow clearance of a large protein alone. First, enhancement persisted through 48 h in PCK kidneys and livers but declined by 48 h in WT organs, and mean enhancement in WT kidneys was about one-third that of PCK kidneys. Second, the nontargeted protein hProCA32, which shares the same scaffold, also enhanced PCK kidneys and livers at early time points but lost 23 to 48% of its peak signal by 48 h, whereas Gd-hProCA32.Collagen retained 88 to 100% (fig. S11 and table S3). This indicated that the protein agent contributes to early enhancement, but that retention depends on the targeting moiety. Third, regional enhancement correlated with CPA in matched histological regions (r = 0.72), whereas Gadovist enhancement, which was small and transient, did not (r = 0.21). Consistent with its pericystic localization, Gd-hProCA32.Collagen increased the number of detected cysts by 25% per animal, compared with 3% for Gadovist, with the largest gains in small and medium cysts. Because each additional cyst is outlined by a collagen-rich matrix, improved delineation of small cysts is a direct consequence of the targeting mechanism.

Current MRI approaches to renal fibrosis, including diffusion-weighted imaging, blood oxygen level–dependent MRI, magnetization transfer, arterial spin labeling, and elastography, provide indirect measures of tissue structure, perfusion, or stiffness that lack specificity for collagen that may be confounded by inflammation, edema, or hemodynamic changes (*35, 36, 94, 95*). Similarly, TKV-based MRI metrics, which remain the clinical standard for ADPKD staging, capture macroscopic cyst burden but fail to detect microcysts and early fibrotic remodeling, resulting in delayed and indirect assessment of disease progression (*3, 34, 96, 97*). In PCK rats, TKV did not distinguish mutant from WT rats at 11 weeks, when renal CPA was already increased. Direct imaging of collagen may therefore complement TKV by reporting a disease process that begins before kidney enlargement becomes measurable.

Several collagen-or matrix-targeted MRI probes have been developed by attaching targeting moieties to small-molecule gadolinium chelates (*98–102*). EP-3533, a collagen-binding peptide conjugated to three Gd-DTPA units, detected myocardial scar and fibrosis in several organs (*103*) but its linear chelators raise concerns about gadolinium release (*104, 105*). CM-101, which carries three macrocyclic Gd-HP-DO3A units, detects liver and lung fibrosis (*106–108*) but renal applications have not been reported. The collagen-binding protein CNA35 and the elastin-specific agent ESMA have also been used for matrix imaging (*109–111*) (*112*) and allysine-targeted probes have been reported that detect oxidized collagen during active fibrogenesis (*113, 114*). Because collagen in early fibrosis is present at low concentrations, sensitivity per gadolinium is a key limitation of these approaches. Gd-hProCA32.Collagen combines high per-gadolinium relaxivity, micromolar affinity for collagen type I preserved in human plasma, and kinetic stability comparable to that of macrocyclic agents. To our knowledge, collagen-targeted MRI has not previously been applied to polycystic kidney disease.

Our regional analyses suggest that cyst expansion and fibrosis are spatially distinct in PCK kidneys: cysts were larger in the OSOM, whereas fibrosis was greater in the cortex, where striation-like enhancement co-localized with clusters of microcysts surrounded by collagen. These observations are consistent with histopathological studies showing that interstitial fibrosis is present early in human ADPKD and in orthologous and nonorthologous rodent models, often coinciding with the earliest cysts (*61, 115*), and that interstitial fibrosis and tubular atrophy are present in human ADPKD kidneys with preserved function (*33, 116*). Fibrotic remodeling of noncystic parenchyma may therefore contribute to nephron loss independently of cyst mass (*6, 117*), and the relationship between cysts and fibrosis is increasingly viewed as bidirectional (*118, 119*). Data from this study do not establish whether fibrosis precedes or drives cyst formation, but they provide a noninvasive means to address this question longitudinally in future studies.

Hepatic cysts and pericystic fibrosis contribute substantially to morbidity in ADPKD but are difficult to quantify noninvasively. Collagen deposition was increased in human polycystic livers and in PCK rat livers, where Gd-hProCA32.Collagen produced sustained enhancement, and hepatic CPA was about threefold higher than in WT rats. Kidney and liver enhancements were also observed in a *Pkd2* mutant mouse model. Because both organs are imaged in the same examination, collagen-targeted MRI could provide a combined assessment of renal and hepatic fibrosis.

Gd-hProCA32.Collagen was administered at a 10-fold lower injection dose than Gadovist, cleared from blood within 72 h, showed transmetallation stability comparable to that of macrocyclic agents, and caused no acute changes in organ weight or histology. These properties are relevant for patients with reduced kidney function, in which gadolinium exposure is a concern. However, residual gadolinium remained in PCK liver and kidney at 72 h, and long-term retention in bone, skin, and brain was not assessed.

The PCK rat model is known for developing progressive hepatorenal disease with preserved renal function during early stages, and this model was used for key pre-clinical studies of tolvaptan. The PCK rat carries a mutation in *Pkhd1*, a gene responsible for autosomal recessive PKD. A conditional floxed *Pkd2* allele in mice was used as a proof-of-concept study. While both the rat and mouse models provided evidence of Gd-hProCA32.Collagen’s utility across cystic diseases, neither directly represents a human patient allele. Larger studies with both *Pkd1* and *Pkd2* conditional mutant mice as well as a mouse with the *Pkd1^RC^*patient variant allele are currently underway. The human specimens analyzed here represent advanced disease, and whether collagen-targeted MRI detects early fibrosis in patients remains to be determined. Since injury accelerates cyst progression and CT contrast agents can induce acute kidney injury responses, the possible deleterious impact of using Gd-hProCA32.Collagen to assess fibrosis needs to be thoroughly evaluated in PKD mutant mouse and rat models prior to translating it to human application.

In conclusion, this work establishes collagen-targeted precision MRI as a transformative approach for early detection and staging of fibrosis in cystic kidney disease, enabling direct visualization of a previously inaccessible subclinical disease phase. By bridging the gap between molecular pathology and noninvasive imaging, hProCA32.Collagen has the potential to complement volumetric imaging for risk stratification and provide a direct readout of fibrosis for trials of antifibrotic therapies in PKD and other chronic kidney diseases.

## Methods and Methods

### Study design

The objective of this study was to develop a collagen-targeted protein MRI contrast agent and to determine whether it detects and maps renal and hepatic fibrosis in polycystic kidney disease. The study comprised (i) quantification of collagen in human ADPKD kidney and polycystic liver tissue; (ii) biophysical characterization of Gd-hProCA32.Collagen (collagen binding, relaxivity, detection limit, serum stability, transmetallation stability, pharmacokinetics, biodistribution, and acute toxicity); (iii) characterization of disease progression in PCK rats by organ weight, MRI-derived total kidney volume (TKV), serum chemistry, and histology; (iv) contrast-enhanced MRI of PCK and WT rats with Gd-hProCA32.Collagen, Gadovist (gadobutrol), or the nontargeted hProCA32, with histological validation; and (v) a proof-of-concept imaging study in *Pkd2* mutant mice. Animals were assigned to contrast agents at random. No animals were excluded. The number of animals and the statistical test for each experiment are given in the figure legends.

### Human tissue

Deidentified formalin-fixed, paraffin-embedded kidney and liver sections were obtained from the UAB Tissue Biorepository (UAB-TBR). They came from patients with ADPKD and from noncystic control tissue. The study was approved by the Institutional Review Board of the University of Alabama at Birmingham (UAB IRB-131223003).

### Contrast agent preparation and characterization

Gd-hProCA32.Collagen and the nontargeted protein hProCA32 were expressed, purified, characterized, and loaded with Gd³⁺ as previously reported (*85, 120*). Protein concentration was determined by Bradford assay and absorbance at 280nm, and Gd content was measured by inductively coupled plasma optical emission spectroscopy (ICP-OES). Unless stated otherwise, doses are expressed as mmol Gd per kg body weight.

### Relaxivity and phantom imaging

Longitudinal and transverse relaxation rates (*R*₁, *R*₂) of Gd-hProCA32.Collagen were measured at 1.4 T (Bruker, minispec) and 9.4 T (Bruker BioSpec) at 37 °C, and per-Gd relaxivities (*r*₁, *r*₂) were calculated from the slopes of *R*₁ and *R*₂ versus [Gd]. Phantoms of Gd-hProCA32.Collagen and Gadovist at the indicated concentrations were imaged at 9.4 T with *T*₁-weighted RARE (TR = 512.82 ms; TE = 12 ms; field of view (FOV) = 63 × 10 mm; matrix = 320 × 128; RARE factor = 2; slice thickness = 0.5 mm; 2 averages) and *T*₂-weighted RARE (TR = 3000 ms; TE = 21.25 ms; RARE factor = 4; other parameters identical). The limit of detection was defined as the lowest concentration producing an SNR increase greater than 3 SD above the buffer baseline.

### Animals

All procedures involving animals were approved by the Institutional Animal Care and Use Committee of the University of Alabama at Birmingham (UAB; protocols #20096 for rats and #21072 for mice) in accordance with the NIH *Guide for the Care and Use of Laboratory Animals*, and with the ARRIVE guidelines. UAB is accredited by AAALAC International.

### PCK rats

Male *Pkhd1*^PCK/PCK^ (PCK) rats, which carry a spontaneous mutation in *Pkhd1* and develop progressive renal and hepatic cysts, were used as a model of polycystic kidney disease (*90, 121*). Male IGS SD rats (Charles River Laboratories) served as wild-type (WT) controls. Rats were studied in the following cohorts: (i) disease characterization: PCK rats at 11, 26, 34, and 41 weeks of age and WT rats at 26 weeks (*n* = 7 ) for serum chemistry, organ weights, and TKV; (ii) contrast-enhanced MRI at 26 weeks: PCK rats receiving Gd-hProCA32.Collagen (*n* = 15), Gadovist (*n* = 9), or hProCA32 (*n* = 3), and WT rats receiving Gd-hProCA32.Collagen or Gadovist (*n* = 5 per agent); (iii) MRI–histology correlation: a subset of PCK rats (*n* = 5 per agent); and (iv) pharmacokinetics, biodistribution, and toxicity: (*n* = 5 per group).

### Pkd2 mice

*Pkd2*^fl/fl^ mice (Jackson Laboratory Stock #017292) were crossed with CaggCre^ER^ (Jackson Laboratory Stock # 004682) mice to establish *Pkd2*^fl/fl^; CaggCre^ER^ conditional mutants. To delete *Pkd2*, mice were injected intraperitoneally with three daily injections of tamoxifen (6mg/40g body weight, Sigma-AldrichCat# T5648-5G) starting at 8 weeks of age. Tamoxifen was dissolved in corn oil, and the mixture was shaken at 37°C until it was dissolved. All mutants and control mice were injected with tamoxifen. A total of 4 mice were studied, comprising 2 Cre^+^ rats and 2 Cre^−^mice.

### MRI acquisition

Imaging was performed at the UAB Small Animal Imaging Facility on a 9.4 T BioSpec scanner (Bruker) with an 86-mm volume coil. Animals were anesthetized with 1.5 to 2% isoflurane, respiration was maintained at 47 to 50 breaths per minute, and body temperature was maintained with a heated bed. Animals were positioned prone, headfirst, and imaged in the axial and coronal planes. The protocol included *T*₂-weighted 2D RARE (TR = 3313 ms; TE = 21.25 ms; FOV = 64 × 64 mm; matrix = 320 × 128; RARE factor = 4; slice thickness = 1 mm; 2 averages); *T*₁-weighted 2D RARE (TR = 1425 ms; TE = 12 ms; other parameters identical); 3D *T*₂-weighted RARE for volumetry (TR = 1000 ms; TE = 80 ms; FOV = 128 × 64 × 64 mm; matrix = 320 × 128 × 64; RARE factor = 8); *T*₁ mapping by FAIR-RARE (TR = 5000 ms; TE = 45.18 ms; RARE factor = 16; inversion times = 50, 300, 600, 1000, 1500, 1800, 2000, and 3000 ms); and *T*₂ mapping by multi-slice multi-echo (TR = 5009 ms; first TE = 9.2 ms).

### Contrast-enhanced imaging

After baseline imaging, Gd-hProCA32.Collagen or hProCA32 (0.010 mmol Gd/kg) or Gadovist (gadobutrol; Bayer HealthCare; 0.1 mmol Gd/kg) was injected into a tail vein at 1 ml/min. Rats were imaged 0.5, 1, 3, 24, and 48 hours after injection. Mice were imaged 1 and 24 hours after injection. For cyst quantification, each agent was analyzed near its time of peak enhancement: 24 hours for Gd-hProCA32.Collagen, which is retained in collagen-rich tissue, and 1 hour for Gadovist, which is cleared from the kidney within hours.

### Ex vivo MRI

After the final *in vivo* scan, kidneys and livers were excised, immersed in phosphate-buffered saline, and imaged at 9.4 T with a 38-mm volume coil using *T*₁-weighted RARE (TR = 938 ms; TE = 12 ms; matrix = 320 × 128; RARE factor = 2; slice thickness = 1 mm; 2 averages) and *T*₂-weighted RARE (TR = 3000 ms; TE = 21.25 ms; RARE factor = 4; other parameters identical).

### Image analysis Signal enhancement

Signal intensities were measured in ImageJ (version 1.53, NIH). Regions of interest (ROIs) were drawn on kidney parenchyma (excluding major vessels, the inner medulla, and the papilla), on individual cysts, on striation regions, and on liver parenchyma, on all slices containing the organ. Post-injection images were matched to pre-injection images using anatomical landmarks. Noise was estimated as the standard deviation of signal in a homogeneous background region outside the animal. SNR was calculated as mean ROI signal divided by noise SD, and percent enhancement as (SNR_post_ − SNR_pre_)/SNR_pre_ × 100. The area under the enhancement–time curve (AUC) from 0 to 48 hours was calculated by the trapezoidal rule (pracma::trapz, R version 4.3.1) and is reported as mean enhancement over the imaging period (AUC/48 hours, %). The retention index was defined as enhancement at 48 hours divided by peak enhancement.

### Total kidney volume

TKV (both kidneys) was measured by manual segmentation of the kidney boundary on *T*₂-weighted images in axial, sagittal, and coronal views in 3D Slicer and Imalytics preclinical software, and calculated from the number of voxels in the resulting mask. *T*₂-weighted images were used for segmentation because cyst fluid is hyperintense and cyst boundaries are well delineated. Measurements were confirmed on *T*₁-weighted images. TKV was normalized to body weight (TKV/BW) and body length (TKV/BL).

### Cyst quantification

Cysts were segmented on *T*₂-weighted images in 3D Slicer by intensity thresholding within the kidney mask, and the number and volume of individual cysts were recorded for both kidneys. The same thresholds were applied to pre-and post-contrast images of each animal. Total cyst volume was summed over both kidneys, and the cyst index was calculated as total cyst volume/TKV × 100. Detection gain was calculated per animal as (post − pre)/pre × 100 for cyst count and total cyst volume. Cysts were classified by volume as micro (0 to 10 mm³), small (11 to 30 mm³), medium (31 to 90 mm³), or large (91 to 150 mm³). Cysts in the cortex and outer stripe of the outer medulla (OSOM) were analyzed separately. Segmentations were repeated twice, and the mean was used (*122, 123*).

### MRI–histology correlation

Five cortical or OSOM ROIs per kidney were defined on post-contrast *T*₂-weighted images in PCK rats (*n* = 5 per agent) and matched to corresponding regions on Picrosirius red–stained sections using anatomical landmarks. Regional percent enhancement was compared with CPA in the matched region.

### Organ weights and serum chemistry

Body weight was recorded at each imaging session. At the end of each study, animals were euthanized under isoflurane anesthesia, and kidneys and liver were excised, blotted, and weighed. Kidney weight is reported for a single kidney (left/right); KW/BW and LW/BW were calculated. Blood was collected from tail veins before contrast administration, and serum was separated by centrifugation (10,000*g*, 10 min, 4 °C) and stored at −20 °C. Serum chemistry (renal, hepatic, electrolyte, metabolic, and enzyme panels) was measured by IDEXX BioAnalytics. Reference ranges are from Charles River Laboratories (*124*).

### Histology and collagen quantification

Tissues were fixed in 10% neutral-buffered formalin overnight, transferred to 70% ethanol, embedded in paraffin, and sectioned at 5 μm. Sections were stained with hematoxylin and eosin, Picrosirius red, and Masson’s trichrome (Abcam, ab150681). Two complementary collagen measures were derived from Picrosirius red–stained sections. (i) Collagen proportionate area (CPA) was quantified in ImageJ as the percentage of Picrosirius red–positive area in 5 non-overlapping fields per section imaged at 4× magnification, and averaged per animal. (ii) A regional fibrosis index was measured on whole digitized sections as described (*122*): Picrosirius red–positive pixels in the cortex and OSOM were detected with a uniform threshold in QuPath and expressed as the fraction of positive area relative to total tissue area in each region. Because the two measures differ in sampling (selected fields versus whole regions) and threshold, their absolute values are not directly comparable.

### Statistical analysis

Analyses were performed in GraphPad Prism (version 11.0.2) and R (version 4.3.1). Data are presented as means ± SD or as box plots showing median, interquartile range, and minimum to maximum, with individual values where shown. The animal was the unit of analysis unless stated otherwise. Two groups were compared with the two-tailed Mann–Whitney test or Welch’s *t-test*, as indicated in the figure legends. Pre-and post-contrast cyst counts and volumes within animals were compared with the Wilcoxon matched-pairs signed rank test. Comparisons of more than two groups used one-way ANOVA with Dunnett’s multiple comparisons test versus WT (serum chemistry) or Kruskal–Wallis with Dunn’s test (CPA at different ages). Correlations were assessed by two-tailed Pearson correlation with linear regression. Because several regions were analyzed per animal, these correlations are reported with the number of regions and animals. Receiver operating characteristic (ROC) analysis used *T*₂-weighted enhancement at 24 h for Gd-hProCA32.Collagen and 1 h for Gadovist; the area under the ROC curve was tested against 0.5. Experiments with *n* = 2 per group (*Pkd2* mice) were analyzed descriptively. *P* < 0.05 was considered significant.

## Data availability

All data associated with this study are present in the paper or the Supplementary Materials. Materials, including Gd-hProCA32.Collagen, are available from J.J.Y. under a material transfer agreement.

## Acknowledgement

We thank B. Rose and R. Guenter (UAB Tissue Biorepository) for assistance with human tissue procurement and T. Du for technical assistance. We thank the UAB Animal Models, Cancer Tissue and Pathology Shared Resources and the UAB Childhood Cystic Kidney Disease Center for assistance in obtaining the *Pkd2* conditional mutant mice, and the UAB Preclinical Imaging Shared Facility for MRI support. We thank Dr. Michael Kirberger for his help in reviewing the manuscript.

## Financial support

This work was supported in part by NIH grants R33CA235319, R42AA112713, R42CA183376, PKD Research Foundation (1560867), and S10OD027045 to J.J.Y., U54DK126087, and R01DK115752. To B.KY. Additional support for commercialization was provided to InLighta BioSciences. Georgia State University CDT fellowship to O.B. and MBD fellowship to F.A. We also acknowledge UAB’s Preclinical Imaging Shared Facility Grant P30CA013148, U54 DK137307, and MRI S10 instrumentation grant S10 OD028498-01.

## Author contributions

Conceptualization: JJY, MM, BKY.

Methodology: JJY, MM, BKY, JW, AGS, KH, AB

Investigation: OB, PHC, FA, JQ, RR, AB, XC, FD, JW

Visualization: OB, AB, JW, KH, AGS

Validation: JJY, MM, BKY, JW

Funding acquisition: JJY, MM, BKY

Supervision: JJY, MM, BKY

Writing original draft: OB, JJY, MM, BKY

## Competing interest

The authors declare the following competing financial interest(s): J.J.Y. holds shares in the company InLighta Biosciences LLC, which licenses the rights to commercialize ProCA agents. J.J.Y. is a named inventor on issued or pending patents (US8173105 (EP1901659), US8367040 (EP3378496) contrast agents, US9339559 (EP1928507, CA 2621763), US 10525150 targeted contrast agents, US9956304 (EP2257316), US15/910893, US17/068215 contrast agents and imaging, US15/572,863 (WO16793465.2) targeted contrast agents. All other authors declare that they have no competing interests.

## Supplementary Materials

Figs. S1 to S14

Table S1 to S3

References

## Notes

### Summary of Updates

Text: The manuscript was edited and condensed throughout to meet journal word limits and style requirements. The Abstract, Introduction, Results, Discussion, and figure legends were rewritten for clarity, and the Discussion now includes an expanded section on study limitations. Figures: The main and supplementary figures were reorganized to meet journal figure limits, and the supplementary figures and tables were renumbered in order of citation. New data: We added imaging data for the nontargeted parental protein hProCA32 as a control (new supplementary figure and table). Analysis: Cyst detection was reanalyzed per animal, comparing the cyst counts and volumes for each rat before and after contrast with Gd-hProCA32.Collagen or gadobutrol (new supplementary table). The correlation between MRI enhancement and histological collagen proportionate area was reanalyzed using matched regions from separate rats for each contrast agent. Statistical tests were revised so that the animal is the unit of analysis, and sample sizes are now reported for every analysis. Methods: The Methods were expanded with details on human tissue procurement and institutional review board approval, contrast agent characterization, image analysis, collagen quantification, and statistics. Other: References, acknowledgments, and funding information were updated.

