## Supplementary Materials for "Unmasking Early Hepatorenal Fibrosis in Polycystic Kidney Disease Using Noninvasive Precision Molecular MRI of Collagen"

Oluwabukola S. Bamishaye, et al.

Corresponding authors:

Jenny J. Yang;

Michal Mrug:

The PDF file includes:

Figs. S1 to S14

Table S1 to S3

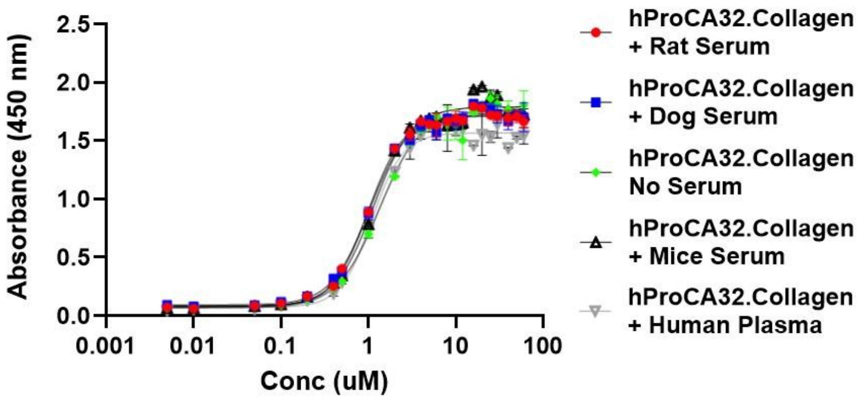


| Best-fit values | hProCA32.Collagen + Rat Serum | hProCA32.Collagen + Dog Serum | hProCA32.Collagen No Serum | hProCA32.Collagen + Mice Serum | hProCA32.Collagen + Human Plasma |
|---|---|---|---|---|---|
| **Bottom** | 0.080 | 0.091 | 0.068 | 0.078 | 0.07180 |
| **Hillslope** | 2.127 | 1.978 | 1.817 | 1.991 | 2.442 |
| **Top** | 1.712 | 1.724 | 1.762 | 1.791 | 1.566 |
| **EC50 (μM)** | **0.998** | **1.029** | **1.336** | **1.145** | **1.092** |
| **logEC50** | -0.001 | 0.012 | 0.126 | 0.059 | 0.038 |
| **Span** | 1.631 | 1.634 | 1.694 | 1.712 | 1.494 |

**Fig. S1. Collagen-binding affinity of hProCA32.Collagen is preserved across serum conditions.** Binding of hProCA32.Collagen to type I collagen was assessed by enzyme-linked immunosorbent assay (ELISA) over a concentration range of 0.001 to 100 μM in the presence of rat, dog, or mouse serum or human plasma, or in the absence of serum. Absorbance was measured at 450 nm. Data are presented as means ± SEM (n = 3 per condition). Curves were fitted using four-parameter nonlinear regression; best-fit parameters are shown in the table. Half-maximal effective concentrations ($EC_{50}$) were 0.998 μM for rat serum, 1.029 μM for dog serum, 1.145 μM for mouse serum, 1.092 μM for human plasma, and 1.336 μM without serum.

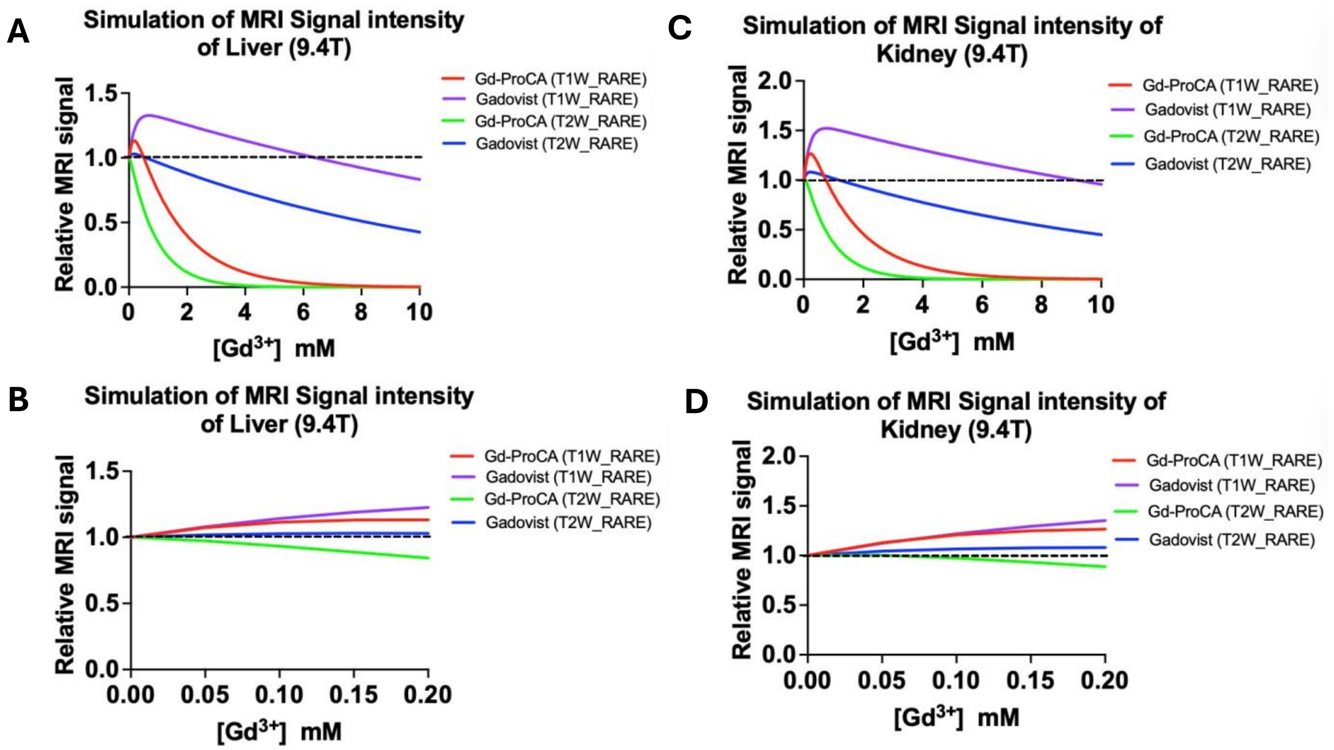


**Fig. S2. Simulated concentration-dependent MRI signal responses to Gd-ProCA and Gadovist in the liver and kidney at 9.4 T.** Relative MRI signal intensity was modeled as a function of gadolinium concentration for Gd-ProCA and Gadovist using T1-weighted (T1W) and T2-weighted (T2W) rapid acquisition with relaxation enhancement (RARE) sequences. (A and B) Simulated liver responses over concentration ranges of 0–10 mM (A) and 0–0.20 mM (B). (C and D) Corresponding kidney responses over concentration ranges of 0–10 mM (C) and 0–0.20 mM (D). Signals were normalized to the pre-contrast baseline (dashed line, relative signal = 1). T1W signal increased at low gadolinium concentrations and decreased at higher concentrations, whereas T2W signal decreased with increasing concentration. Red and purple curves indicate T1W responses for Gd-ProCA and Gadovist, respectively; green and blue curves indicate their corresponding T2W responses. These simulations inform optimal sequence selection for detecting collagen-targeted contrast enhancement *in vivo* at high field strength.

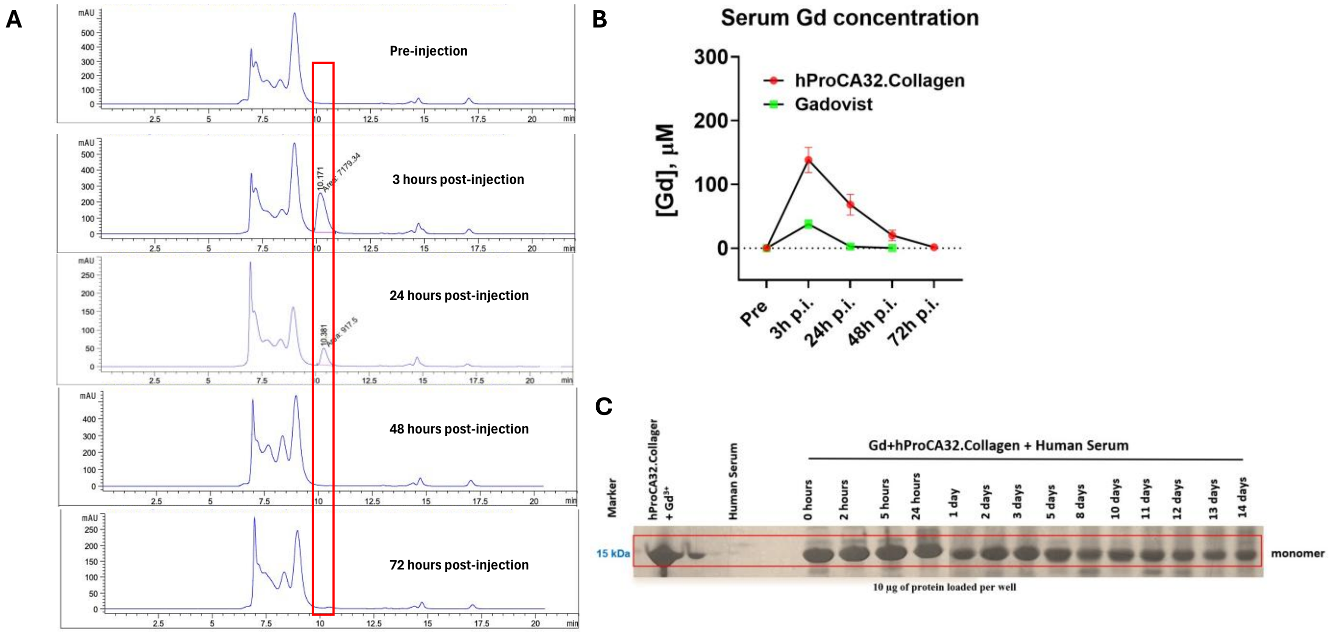


**Fig. S3. Blood clearance and serum stability of Gd-hProCA32.Collagen.** (A) Size-exclusion high-performance liquid chromatography (SEC-HPLC) profiles of rat serum collected before and at 3, 24, 48, and 72 hours after intravenous administration of Gd-loaded hProCA32.Collagen (0.010 mmol Gd/kg). The red box indicates the elution region corresponding to intact Gd-loaded hProCA32.Collagen (retention time, approximately 10.17 min). (B) Total serum gadolinium concentrations before and after intravenous administration of Gd-loaded hProCA32.Collagen (0.010 mmol Gd/kg) or Gadovist (gadobutrol; 0.100 mmol Gd/kg). Data are means ± SD; *n* = 5 rats per group. (C) Coomassie-stained SDS–polyacrylamide gel showing Gd-loaded hProCA32.Collagen after incubation in human serum at 37°C for up to 14 days. Control lanes contain a molecular weight marker, Gd-loaded hProCA32.Collagen, and human serum alone. The red box highlights the monomer band at approximately 15 kDa. Protein loading was 10 µg per lane.

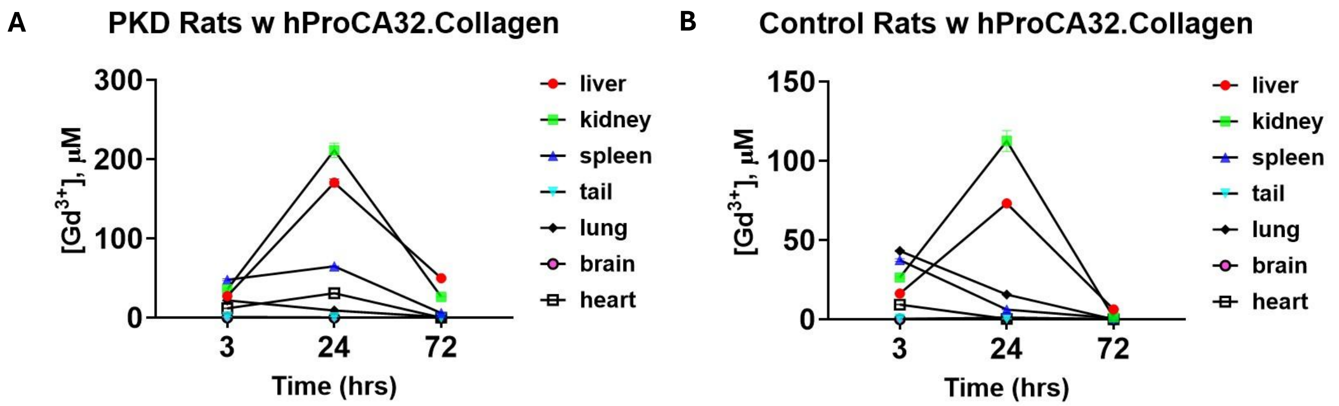


**Fig. S4. Tissue distribution of gadolinium in PKD and control rats after administration of hProCA32.Collagen.** Gadolinium concentrations in the liver, kidney, spleen, tail, lung, brain, and heart were measured at 3, 24, and 72 hours after administration of Gd-loaded hProCA32.Collagen in (A) rats with polycystic kidney disease (PKD) and (B) control rats. In both groups, liver and kidney gadolinium concentrations were highest at 24 hours among the sampled time points and decreased by 72 hours. Minimal brain accumulation was observed in both groups, indicating limited blood–brain barrier penetration. Data demonstrate preferential accumulation and prolonged retention of hProCA32.Collagen in fibrotic PKD-associated organs, with subsequent systemic clearance over time. Gd-hProCA32.Collagen (0.010 mmol Gd/kg) was injected intravenously, and Gd concentration was measured using ICP-OES; n=5 in each group.

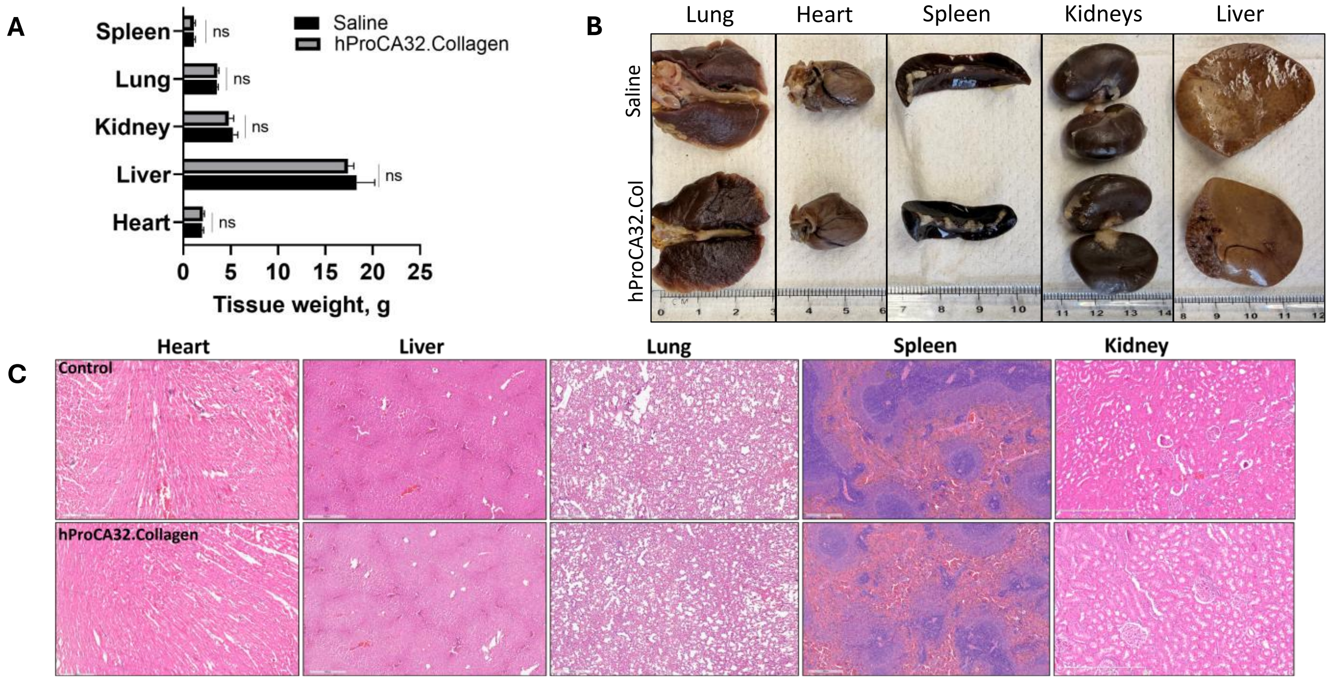


**Fig. S5. Organ weights, gross morphology, and histology in rats after Gd-hProCA32.Collagen administration.** Rats received intravenous injections of Gd-hProCA32.Collagen (0.010 mmol Gd/kg) or saline, and organs were collected 4 days after injection ($n$ = 5 rats per group). (A) Heart, liver, kidney, lung, and spleen weights in saline-treated and Gd-hProCA32.Collagen-treated rats. Data are means ± SEM. Statistical comparisons were performed using a *t-test*; ns, not significant ($P > 0.05$). (B) Representative gross images of the lung, heart, spleen, kidneys, and liver from saline injected (top row) and Gd-hProCA32.Collagen injected (bottom row) rats. Rulers provide size references. (C) Representative hematoxylin and eosin–stained sections of the heart, liver, lung, spleen, and kidney from saline-treated (top row) and Gd-hProCA32.Collagen-treated (bottom row) rats. Scale bars, 600 µm.

**Table S1**. Serum chemistry in WT and PCK rats. Serum was collected from WT rats at 26 weeks and PCK rats at 11, 26, 34, and 41 weeks of age (before contrast agent administration). Values are means ± SD; n = 7 rats per group. Reference ranges (mean) are for Sprague-Dawley rats from Charles River Laboratories (*1*).

| | **WT Rat** | **PCK 11 Weeks** | **PCK 26 Weeks** | **PCK 34 Weeks** | **PCK 41 Weeks** | **REFERENCE (RANGE) MEAN** |
|---|---|---|---|---|---|---|
| **Creatine Kinase (U/L)** | 1328 | 1131 | 1009 | 780.5 | 211.0 | (%6–477) 241 |
| **Amylase (U/L)** | 662.0 | 847.0 | 781.7 | 821.0 | - | (500–1200) 850 |
| **Glucose (mg/dL)** | 414.3 | 123.5 | 538.0 | 290.0 | 158.5 | (92–138) 115 |
| **LDH (U/L)** | 444.3 | 366.0 | 364.7 | 249.0 | - | (76–953) 210 |
| **GGT (U/L)** | 0.000 | 0.333 | 1.000 | 1.000 | - | (0–3) 0.5 |
| **Albumin (g/dL)** | 3.400 | 2.950 | 2.533 | 2.450 | 2.150 | (3.7–5.0) 4.45 |
| **Total Bilirubin (mg/dL)** | 0.100 | 0.100 | 0.100 | 0.150 | 0.700 | (0.10–0.20) 0.15 |
| **Total Protein (g/dL)** | 7.333 | 5.750 | 6.333 | 6.400 | 6.950 | (6.3–8.1) 6.7 |
| **Globulin (g/dL)** | 3.933 | 2.800 | 3.800 | 3.950 | 4.800 | (1.5–3.0) 2.2 |
| **Creatinine (mg/dL)** | 0.300 | 0.373 | 0.713 | 0.585 | 0.230 | (0.4–0.6) 0.57 |
| **Phosphorus (mg/dL)** | 9.900 | 8.950 | 8.733 | 7.200 | 4.900 | (5.50–9.20) 7.65 |
| **Potassium (mmol/L)** | 8.433 | 8.333 | 8.633 | 8.050 | 4.800 | (4.2–6.7) 5.65 |
| **ALB/GLOB Ratio** | 0.867 | 1.050 | 0.667 | 0.600 | 0.500 | (0.8–2.0) 1.2 |
| **Bilirubin (mg/dL)** | 0.100 | 0.100 | 0.100 | 0.100 | 0.600 | (0.0–0.15) 0.08 |
| **Uric Acid (mg/dL)** | 8.533 | 8.333 | 9.033 | 7.250 | 12.00 | (1.2–7.5) 3.5 |
| **ALP (U/L)** | 131.0 | 141.5 | 171.0 | 150.0 | 239.5 | (53–226) 105 |
| **AST (U/L)** | 120.7 | 86.00 | 118.3 | 83.50 | 169.0 | (57–144) 90.5 |
| **ALT (U/L)** | 62.67 | 37.50 | 58.33 | 79.50 | 80.00 | (26–97) 47 |
| **Lipase (U/L)** | 31.33 | 64.33 | 77.00 | 82.50 | - | (20–160) 60 |
| **BUN (mg/dL)** | 18.67 | 16.00 | 23.67 | 35.00 | 16.00 | (15–80) 40 |
| **Cholesterol (mg/dL)** | 97.33 | 100.0 | 132.3 | 157.5 | 73.50 | (62–234) 92 |
| **Calcium (mg/dL)** | 12.90 | 8.567 | 12.13 | 12.45 | 9.900 | (9.9–11.9) 10.9 |
| **Bicarbonate TCO2 (mmol/L)** | 26.00 | 22.00 | 24.33 | 25.00 | 31.00 | (18–28) 23 |
| **Chloride (mmol/L)** | 94.67 | 92.00 | 89.67 | 100.5 | 100.5 | (95–111) 103.5 |
| **Sodium (mmol/L)** | 143.7 | 144.0 | 138.0 | 144.5 | 140.5 | (143–157) 148 |
| **BUN/Creatinine Ratio** | 62.97 | 48.40 | 37.87 | 60.00 | 70.20 | (15–80) 40 |
| **NA/K Ratio** | 17.00 | 14.33 | 12.33 | 18.00 | 29.00 | (15–30) 22 |
| **Triglycerides (mg/dL)** | 226.7 | 217.7 | 197.3 | 236.5 | - | (46–208) 94 |

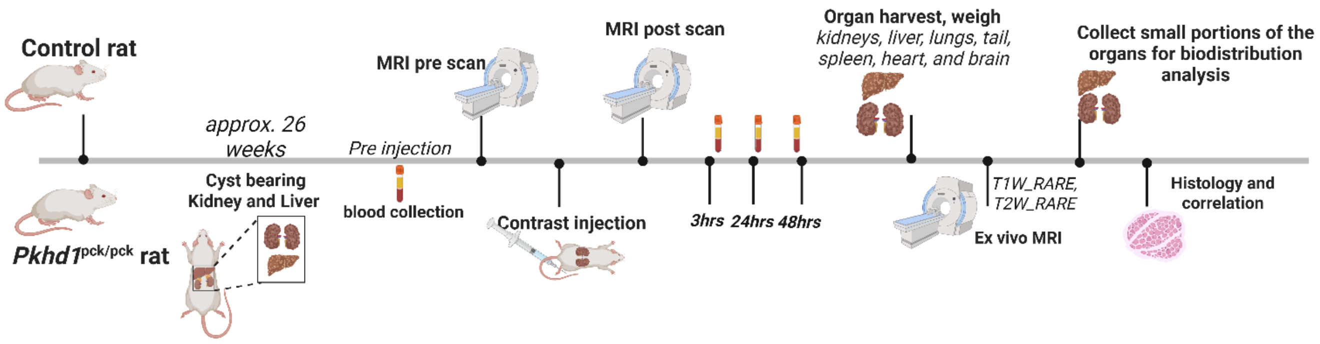


**Fig. S6. Experimental design for *in vivo* MRI and tissue analyses.** WT and *Pkhd1*$^{PCK/PCK}$ (PCK) rats (26 weeks old) underwent blood collection and pre-contrast MRI ($T_1$- and $T_2$-weighted RARE imaging and $T_1$ and $T_2$ mapping), followed by intravenous injection of contrast agent and post-contrast MRI. Blood was collected 3, 24, and 48 hours after injection. Kidneys, liver, lungs, spleen, heart, brain, and tail were then harvested and weighed; excised kidneys and liver were imaged *ex vivo*, and tissue was collected for biodistribution and histological analysis. *Created in BioRender. Yang lab, J. (2026)* https://BioRender.com/z4ey04m

**Table S2. Per-animal cyst detection in PCK rats imaged with Gadovist or Gd-hProCA32.Collagen.**

| | Gd-hProCA32.Collagen ($n$ = 15) | Gadovist ($n$ = 9) | $P$ |
|---|---|---|---|
| ***Baseline*** | | | |
| Cyst count | 70.9 ± 16.4 | 78.9 ± 27.1 | 0.44 |
| Total cyst volume ($mm^3$) | 1,186 ± 450 | 1,425 ± 666 | 0.36 |
| Total kidney volume ($mm^3$) | 8,571 ± 1,947 | 9,435 ± 2,714 | 0.42 |
| Cyst index (%) | 13.8 ± 3.4 | 15.5 ± 6.5 | 0.48 |
| ***After contrast*** | | | |
| Additional cysts detected | 17.0 ± 3.9 | 1.8 ± 1.2 | <0.0001 |
| Increase in cyst count (%) | 25.1 ± 8.4 | 2.6 ± 2.0 | <0.0001 |
| Increase in cyst volume (%) | 16.3 ± 10.1 | 0.7 ± 0.8 | <0.0001 |

*Values are means ± SD. Gd-hProCA32.Collagen (0.010 mmol Gd/kg) was imaged 24 hours and Gadovist (gadobutrol, 0.1 mmol Gd/kg) 1 hour after intravenous injection on $T_2$-weighted MRI in 26-week-old PCK rats. Cyst index = (total cyst volume/TKV) × 100. Percent increases were calculated per animal as (post − pre)/pre × 100. Baseline values were compared with Welch's t-test and post-contrast changes with the two-tailed Mann–Whitney test.*

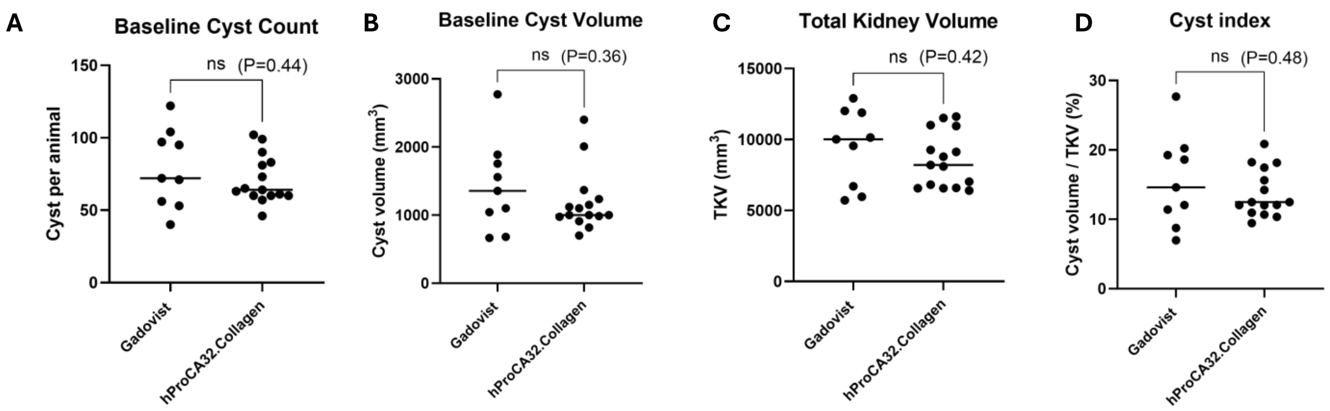


**Fig. S7. Baseline renal disease burden in PCK rats assigned to Gadovist or Gd-hProCA32.Collagen imaging.** (A) Cyst count, (B) total cyst volume, (C) total kidney volume (TKV), and (D) cyst index (total cyst volume/TKV × 100) on pre-contrast $T_2$-weighted MRI in 26-week-old PCK rats subsequently imaged with Gadovist (gadobutrol; n = 9) or Gd-hProCA32.Collagen (n = 15). Each dot represents one rat; horizontal lines indicate means. Welch's t-test; P = 0.44, 0.36, 0.42, and 0.48 for (A) to (D), respectively. ns, not significant.

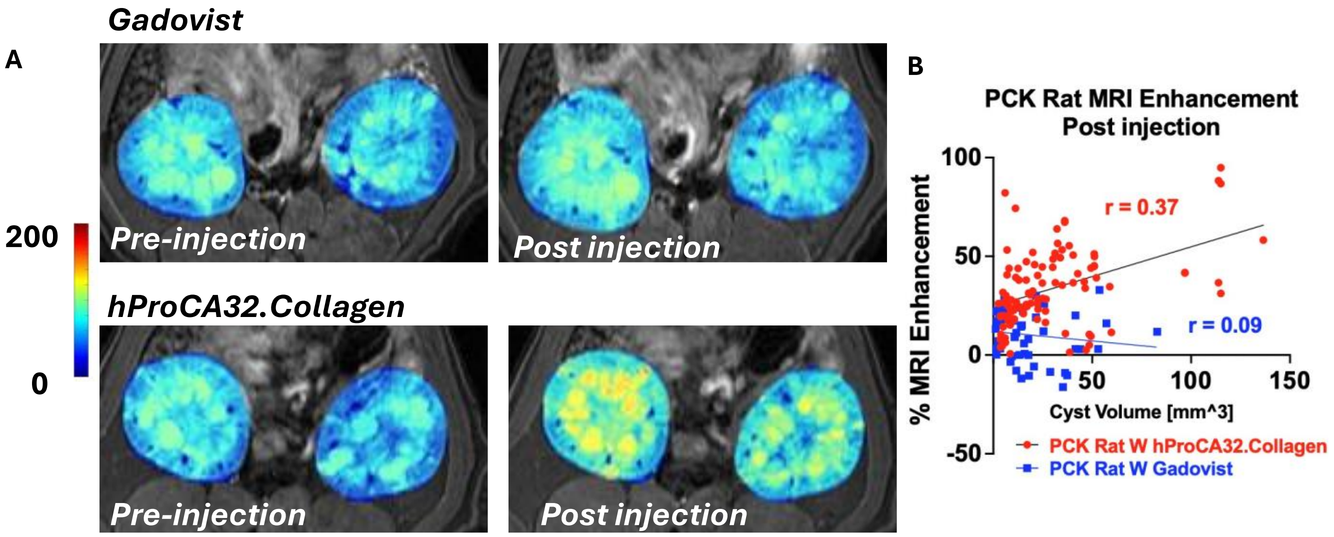


**Fig. S8. Renal signal maps and relationship between enhancement and cyst volume in PCK rats**. (A) Representative $T_2$-weighted SNR maps of PCK rat kidneys before and after Gadovist (0.1 mmol Gd/kg; top) or Gd-hProCA32.Collagen (0.010 mmol Gd/kg; bottom). (B) Percent signal enhancement plotted against volume for individual cysts after Gd-hProCA32.Collagen (red circles; $r = 0.37$) or Gadovist (blue squares; $r = 0.09$), with linear regression lines. Cysts were pooled from $n = 15$ (Gd-hProCA32.Collagen) and $n = 9$ (Gadovist) rats.

s

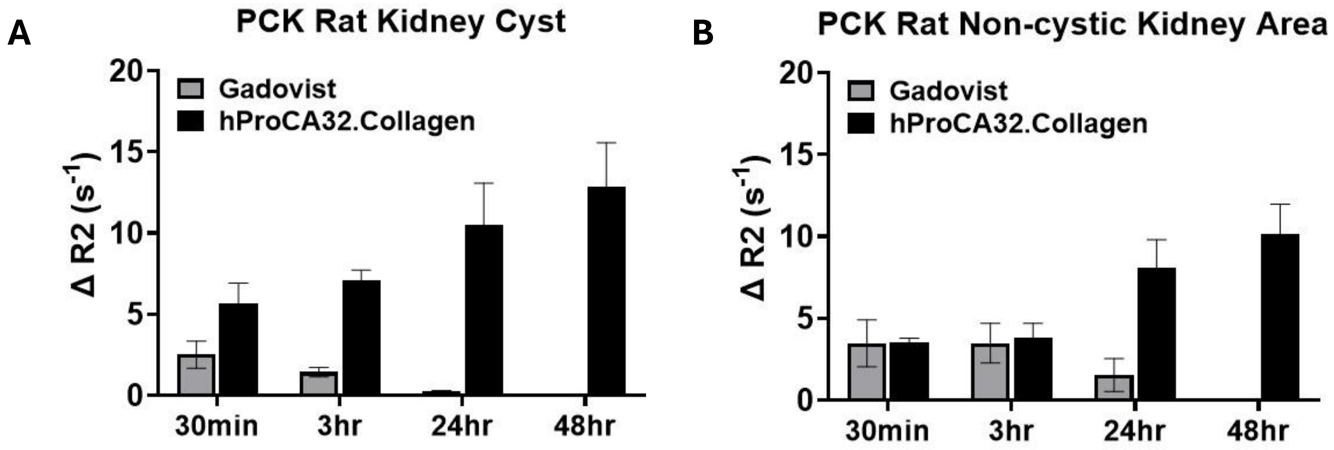


**Fig. S9. Transverse relaxation rate changes in cystic and noncystic kidney regions of PCK rats.** Change in transverse relaxation rate from baseline ($\Delta R_2$, $s^{-1}$) in (A) kidney cysts and (B) noncystic renal parenchyma at 30 min and 3, 24, and 48 hours after Gd-hProCA32.Collagen (0.010 mmol Gd/kg; black bars) or Gadovist (0.1 mmol Gd/kg; gray bars). Gadovist was not measured at 48 hours. Data are means ± SD; *n* = 15 (Gd-hProCA32.Collagen) and 9 (Gadovist) rats.

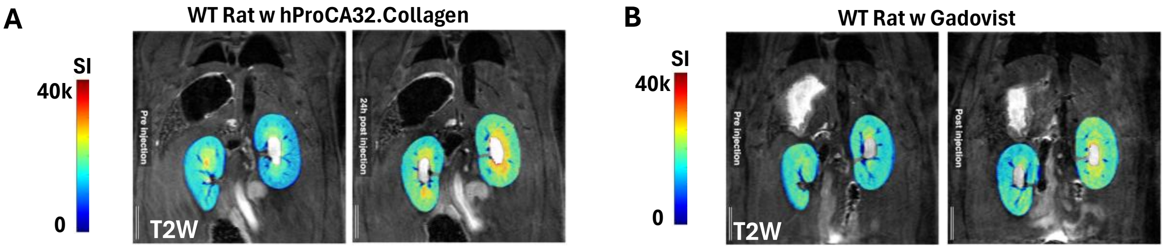


**Fig. S10. $T_2$-weighted signal intensity maps of WT rat kidneys before and after contrast.** Representative $T_2$-weighted images with renal signal intensity (SI) overlays before and (A) 24 hours after Gd-hProCA32.Collagen or (B) 1 hour after Gadovist in WT rats. Color scale, SI 0 (blue) to 40,000 (red). $n$ = 5 rats per agent.

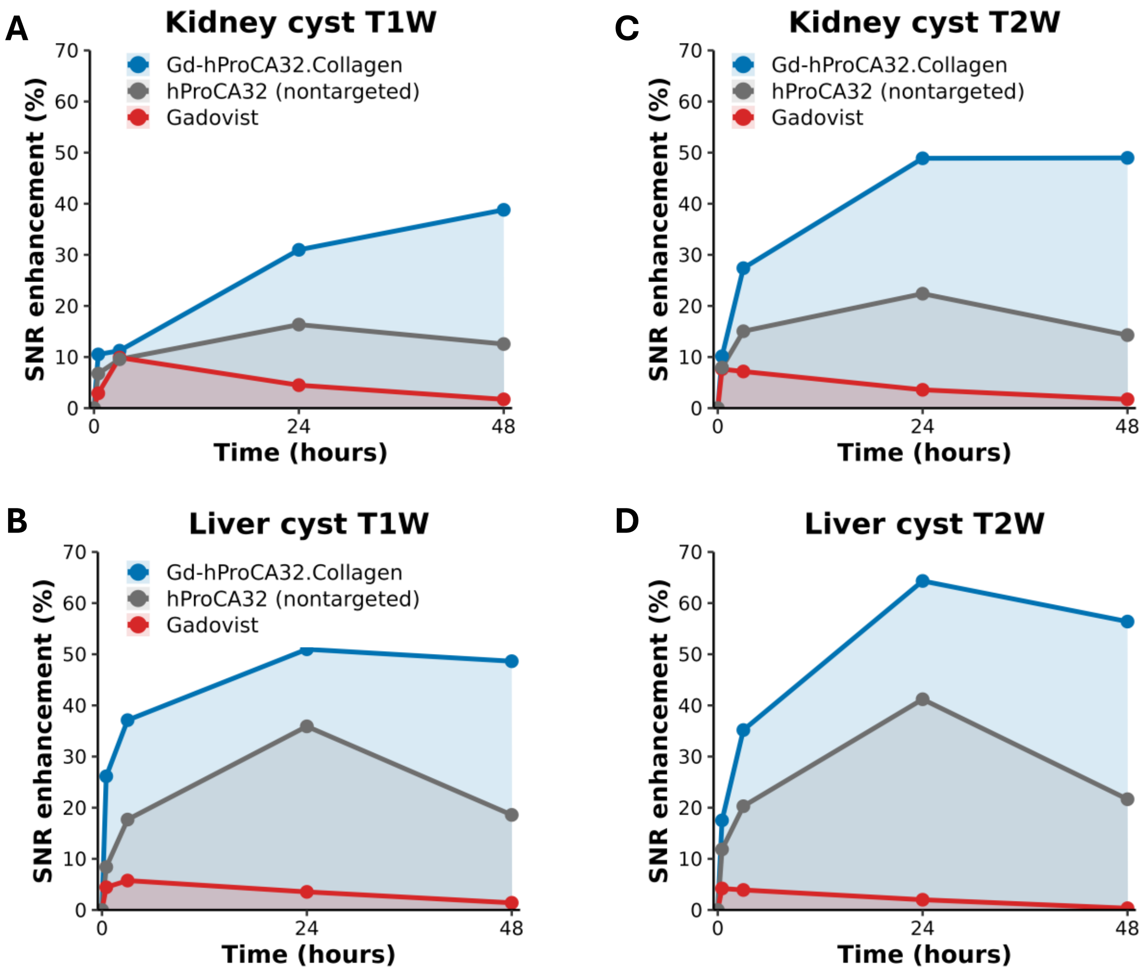


**Fig. S11. Sustained renal and hepatic enhancement depends on collagen targeting.** MRI SNR enhancement (%) over 48 hours after intravenous injection of Gd-hProCA32.Collagen (blue; 0.010 mmol Gd/kg), the nontargeted parental protein hProCA32 (gray; 0.010 mmol Gd/kg), or Gadovist (red; 0.1 mmol Gd/kg) in 26-week-old PCK rats. (A and B) T1-weighted enhancement of kidney cysts (A) and liver cysts (B). (C and D) T2-weighted enhancement of kidney cysts (C) and liver cysts (D). Images were acquired before and 0.5, 3, 24, and 48 hours after injection. Shaded areas show the area under the curve. Data are group means *n* = 15 (Gd-hProCA32.Collagen), 9 (Gadovist), and 3 (hProCA32) rats. Kinetic parameters are summarized in table S3.

**Table S3. Kidney and liver enhancement kinetics in PCK rats after collagen-targeted, nontargeted, and clinical contrast agents.**

| Organ | Sequence | Contrast agent | Peak enhancement, % (time) | Enhancement at 48 h (%) | Retention index (48 h/peak) | $AUC_{0–48\ h}$ (%·h) | Mean enhancement, 0–48 h (%) |
|---|---|---|---|---|---|---|---|
| Kidney | T1W | Gd-hProCA32.Collagen | 38.8 ± 1.2 (48 h) | 38.8 ± 1.2 | 1.00 | 1,311 | 27.3 ± 1.4 |
| | | hProCA32 | 16.3 ± 0.8 (24 h) | 12.5 ± 0.8 | 0.77 | 641.0 | 13.4 ± 0.6 |
| | | Gadovist | 9.90 ± 0.4 (3 h) | 1.70 ± 0.4 | 0.17 | 242.0 | 5.10 ± 0.2 |
| | T2W | Gd-hProCA32.Collagen | 49.0 ± 2.0 (48 h) | 49.0 ± 2.0 | 1.00 | 2,025 | 41.6 ± 0.8 |
| | | hProCA32 | 22.4 ± 1.3 (24 h) | 14.3 ± 1.1 | 0.64 | 864.0 | 18.0 ± 0.3 |
| | | Gadovist | 7.60 ± 0.4 (0.5 h) | 1.70 ± 0.3 | 0.22 | 197.0 | 4.20 ± 0.1 |
| Liver | T1W | Gd-hProCA32.Collagen | 51.0 ± 2.3 (24 h) | 48.6 ± 1.8 | 0.95 | 2,206 | 46.0 ± 0.2 |
| | | hProCA32 | 35.9 ± 1.8 (24 h) | 18.6 ± 0.7 | 0.52 | 1,251 | 26.1 ± 0.2 |
| | | Gadovist | 5.80 ± 1.3 (3 h) | 1.40 ± 0.3 | 0.24 | 171.0 | 3.60 ± 0.1 |
| | T2W | Gd-hProCA32.Collagen | 64.3 ± 3.1 (24 h) | 56.4 ± 3.0 | 0.88 | 2,564 | 53.4 ± 0.2 |
| | | hProCA32 | 41.2 ± 2.1 (24 h) | 21.7 ± 0.6 | 0.53 | 1,443 | 30.1 ± 0.1 |
| | | Gadovist | 4.20 ± 1.7 (0.5 h) | 0.30 ± 0.01 | 0.08 | 101.0 | 2.10 ± 0.1 |

*Values are group means of percent SNR enhancement in PCK rats imaged at 0.5, 3, 24, and 48 hours after intravenous injection of Gd-hProCA32.Collagen (0.010 mmol Gd/kg), nontargeted hProCA32 (0.010 mmol Gd/kg), or Gadovist (0.1 mmol Gd/kg); n = 15 for rats Gd-hProCA32.Collagen, 9 (Gadovist), and 3 (hProCA32) rats. AUC was calculated by the trapezoidal rule from 0 to 48 hours; mean enhancement = AUC/48 hours. Retention index = enhancement at 48 hours divided by peak enhancement. SNR, signal-to-noise ratio; T1W and T2W, T1- and T2-weighted RARE imaging*

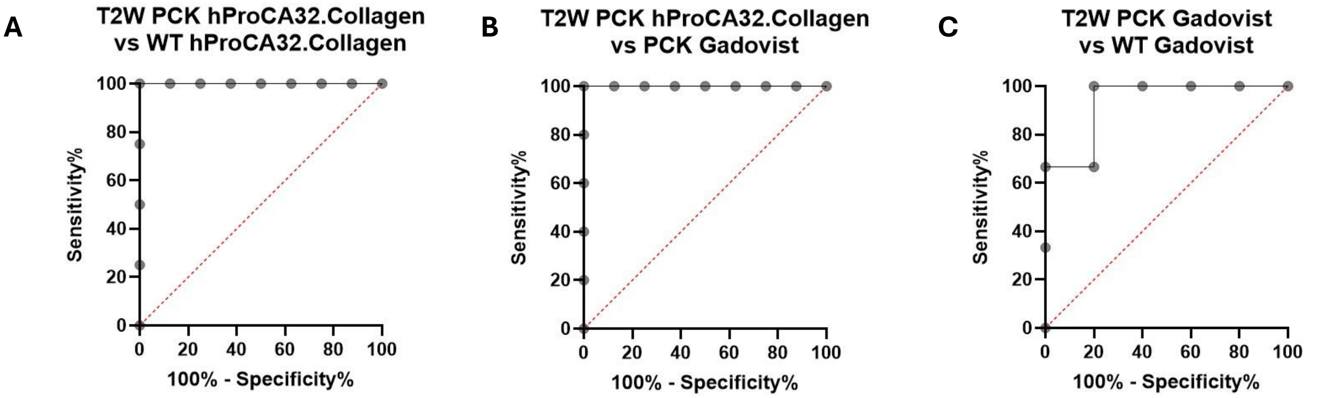


| Area under the ROC curve | |
|---|---|
| Area | 1.000 |
| Std. Error | 0.000 |
| 95% confidence interval | 1.000 to 1.000 |
| P value | 0.0066 |

| Area under the ROC curve | |
|---|---|
| Area | 1.000 |
| Std. Error | 0.000 |
| 95% confidence interval | 1.000 to 1.000 |
| P value | 0.0034 |

| Area under the ROC curve | |
|---|---|
| Area | 0.9333 |
| Std. Error | 0.09428 |
| 95% confidence interval | 0.7485 to 1.000 |
| P value | 0.0526 |

**Fig. S12. Receiver operating characteristic analysis of $T_2$-weighted MRI enhancement.** ROC curves of $T_2$-weighted SNR enhancement measured 24 hours after Gd-hProCA32.Collagen or 1 hour after Gadovist. (A) PCK ($n$ = 15) versus WT ($n$ = 5) rats receiving Gd-hProCA32.Collagen: area under the curve (AUC), 1.00; 95% confidence interval (CI), 1.00 to 1.00; $P$ = 0.007. (B) PCK rats receiving Gd-hProCA32.Collagen ($n$ = 15) versus Gadovist ($n$ = 9): AUC, 1.00; 95% CI, 1.00 to 1.00; $P$ = 0.003. (C) PCK ($n$ = 9) versus WT ($n$ = 5) rats receiving Gadovist: AUC, 0.93; 95% CI, 0.75 to 1.00; $P$ = 0.053. Dashed lines indicate chance discrimination (AUC = 0.5).

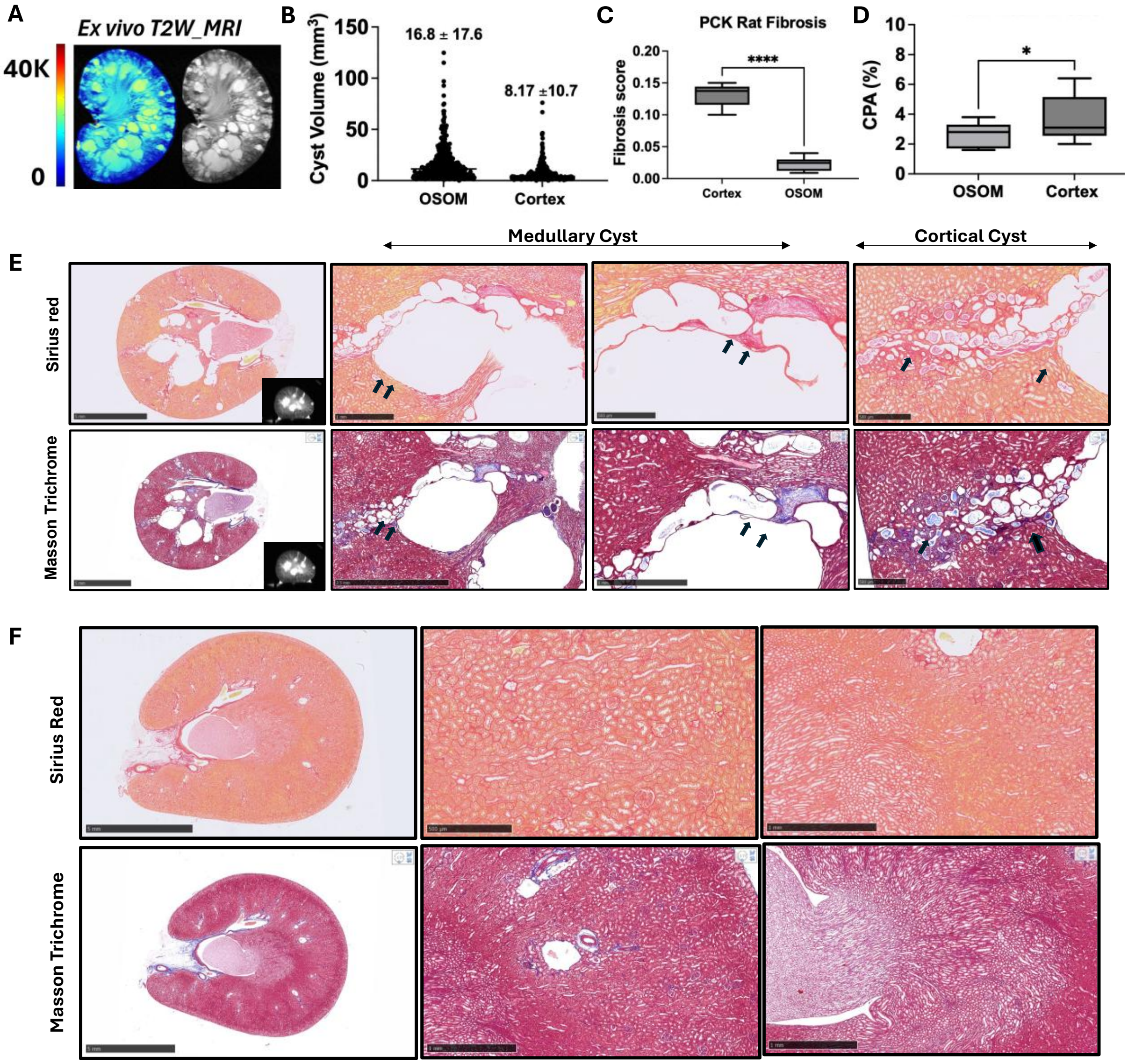


**Fig. S13. Regional distribution of cysts and fibrosis in PCK rat kidneys**. (A) *Ex vivo* $T_2$-weighted signal intensity map (left) and grayscale image (right) of an excised 26-week-old PCK kidney. (B) Volumes of individual cysts in the outer stripe of the outer medulla (OSOM) and cortex measured on MRI; each point is one cyst from $n = 10$ rats; values are means ± SD. (C) Regional fibrosis index (fraction of Picrosirius red–positive area in whole cortical and OSOM regions), measured as described (*2*). (D) Collagen proportionate area (CPA; percentage of Picrosirius red–positive area in selected fields) in the OSOM and cortex. (E) Representative Picrosirius red-stained (top) and Masson's trichrome-stained (bottom) PCK kidney sections, with *ex vivo* MRI insets and higher-magnification views of medullary and cortical cysts; arrows indicate pericystic and interstitial collagen. (F) Representative Picrosirius red–stained (top) and Masson's trichrome–stained (bottom) WT kidney sections at low and high magnification. Scale bars are shown. $n = 10$ rats in (C) and (D); Mann–Whitney test. $^{*}P < 0.05$, $^{****}P < 0.0001$.

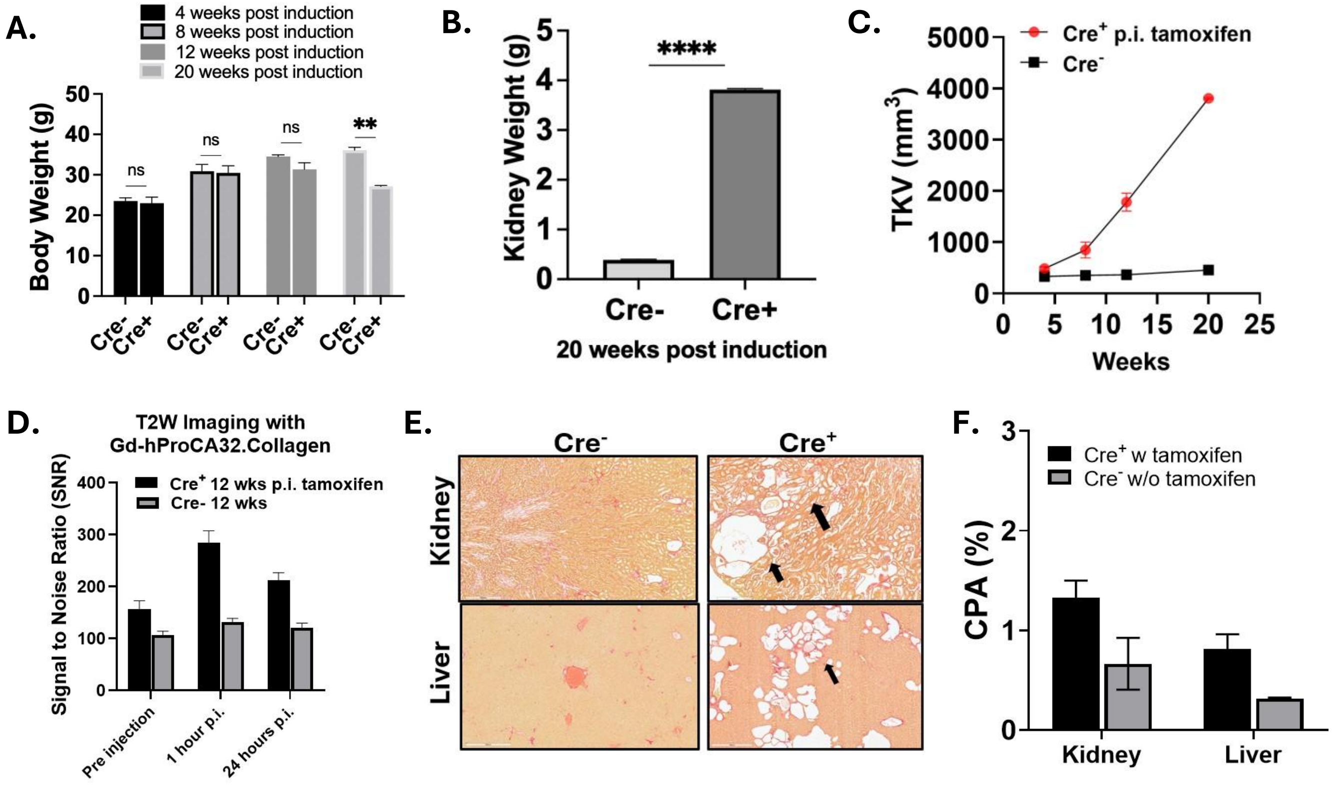


**Fig. S14. Disease progression and collagen-targeted MRI in tamoxifen-inducible *Pkd2* mutant mice.** Male mice were induced with tamoxifen at 4 weeks of age; uninduced $Cre^-$ littermates served as controls. (A to C) Longitudinal cohort: body weight at 4, 8, 12, and 20 weeks after induction (A), kidney weight (both kidneys) at 20 weeks after induction (B), and MRI-derived total kidney volume (TKV) from 4 to 20 weeks after induction (C). (D) $T_2$-weighted kidney SNR before and 1 and 24 hours after Gd-hProCA32.Collagen (0.010 mmol Gd/kg) 12 weeks after induction. (E) Representative Picrosirius red stained kidney (top) and liver (bottom) sections from $Cre^-$ (left) and $Cre^+$ (right) mice collected 24 hours after Gd-hProCA32.Collagen injection, 12 weeks after induction; arrows indicate pericystic collagen. (F) CPA in kidney and liver of the imaged mice. Data are means ± SD (A to C) or means with individual values (D and F); $n = 2$ per group in (A) to (C) and 2 per group in (D) to (F). Welch's *t-test* (A and B); $^{**}P < 0.01$, $^{****}P < 0.0001$; ns, not significant. Scale bars are shown.
